# Bayesian Modeling of Context-Specific Goal-Directed and Habitual Control in Sequential Decision Making

**DOI:** 10.64898/2026.11.06.764848

**Authors:** Willi Hieke, Sarah Schwöbel, Sia Hranova, Arvid Pietsch, Stefan J. Kiebel, Michael N. Smolka

## Abstract

It is an open question how context-specific habits develop alongside goal-directed planning. We examined action sequence repetition, contingency change, and context switching using a three-step optimization task with interleaved planning and habit contexts. During an initial acquisition phase, optimal sequences varied in the planning context but remained constant in the habit context. When contingencies changed, the original habit sequence became suboptimal and another became optimal. Action sequence repetition improved response speed and optimal-choice performance in the habit context relative to the planning context. Following the contingency change, both advantages initially reversed. Optimal-choice performance in the habit context subsequently recovered but did not significantly exceed that in the planning context, while responses remained slower in the habit context. Context switches reduced optimal-choice performance in both directions, with additional slowing when switching from habit to planning. The Bayesian contextual control model captured choice patterns through learned action sequence priors, goal-directed evaluation, and context inference. Allowing a context-stability bias to vary across participants did not improve model performance. These findings suggest that sequence repetition facilitates performance under stable contingencies while impairing adaptation to change.

## 1 Introduction

The study of habitual and goal-directed behavior has a long tradition in psychology and neuroscience. Classical accounts distinguish between goal-directed actions, which depend on action-outcome contingencies and current outcome values, and habitual actions, which are relatively insensitive to such changes and are often described as stimulus-response associations [Dickinson, 1985, Balleine and Dickinson, 1998]. Subsequent work in animals linked these behavioral systems to distinct neural substrates, particularly within cortico-striatal circuits, with the dorsomedial striatum implicated in goal-directed control and the dorsolateral striatum in habitual responding [Yin et al., 2004, Yin and Knowlton, 2006, Graybiel, 2008, Dolan and Dayan, 2013].

In cognitive neuroscience, this distinction has often been interpreted through the lens of reinforcement learning (RL). In particular, goal-directed behavior has been formalized as model-based RL, which relies on an internal model of state transitions, whereas habitual behavior has been formalized as model-free RL, which relies on cached value estimates [Daw et al., 2005, Daw et al., 2011, Dolan and Dayan, 2013]. This framework has been supported by a large body of behavioral and neuroimaging research, most prominently using the two-step task, which dissociate model-based and model-free contributions to choice behavior [Daw et al., 2011, Gläscher et al., 2010]. The two-step task has also been used to investigate differences between healthy individuals and those with mental disorders [Gillan et al., 2016], or risky substance use [Chen et al., 2021, Chen et al., 2026]. Related work has further examined how control may be allocated between these two modes, for example through arbitration based on uncertainty or reliability estimates, and has highlighted the contribution of perseveration or repetition biases to sequential choice behavior [Lee et al., 2014, Akam et al., 2015, Kool et al., 2016].

The identification of habits with model-free RL, and more generally the strict dichotomy between habitual and goal-directed behavior, remains debated (see, e.g., [Friedel et al., 2014]). One alternative is provided by value-free stimulus–response accounts, which propose that habits may form through the direct strengthening of action tendencies by repetition, independently of outcome value [Miller et al., 2019]. This view is consistent with recent empirical findings from studies in humans and animals [Frölich et al., 2023, Greenstreet et al., 2025, Lebedeva et al., 2026].

Furthermore, computational frameworks such as the *successor representation* combine features of model-based and model-free control, suggesting that behavioral flexibility may not be fully captured by a simple two-system dichotomy [Dayan, 1993, Gershman, 2018]. In parallel, habits have been proposed to operate at the level of action sequences rather than individual actions, with repeated actions becoming chunked or hierarchically organized in sequential decision making tasks [Dezfouli and Balleine, 2012, Dezfouli et al., 2014, Smith and Graybiel, 2016, Frölich et al., 2023, Grundmann et al., 2025].

An alternative line of work, based on the *Bayesian brain hypothesis* and the *active inference* framework, conceptualizes behavior as probabilistic inference over hidden states and action sequences [Knill and Pouget, 2004, Friston, 2010, Friston et al., 2015]. Within this perspective, the Bayesian contextual control (BCC) model [Schwöbel et al., 2021] combines both value-free habit learning and chunked, hierarchically organized action sequences. Previous work has long emphasized that habits are context-specific [Wood and Rünger, 2016]. The BCC model incorporates this idea by assuming that action selection is guided by beliefs about the current context, where goal-directed behavior is guided by expected outcomes, whereas habitual behavior reflects the influence of learned priors over actions or action sequences. Accordingly, the balance between habitual and goal-directed behavior follows from inference over contexts and action sequences.

Previous empirical studies of habitual behavior in humans have predominantly relied on outcome-devaluation and contingency-degradation paradigms [Yin and Knowlton, 2006, Gillan et al., 2015]. These paradigms have been instrumental in establishing key characteristics of habit learning, including reduced sensitivity to changes in outcome value and action–outcome contingencies, as well as effects of training duration on the strength of habitual responding [Yin and Knowlton, 2006, Seger and Spiering, 2011].

However, existing habit paradigms often involve either limited sequential structure or do not explicitly incorporate context. These limitations are important because realworld habits often unfold over extended action sequences whose expression depends on context [Lally et al., 2010, Wood and Rünger, 2016, Verplanken and Roy, 2016]. Previous work has shown that habit acquisition depends on context consistency and that changes in context can disrupt habitual responding [Danner et al., 2008, Lally et al., 2010, Neal et al., 2012]. These findings motivate paradigms that capture both the sequential and context-dependent nature of human habit learning. Moreover, a recent meta-analysis in the clinical domain of substance use disorder (SUD) highlights the importance of translating the habit construct across experimental, computational, and applied settings [Giannone et al., 2024]. Studies on SUD have reported differences in learning and reliance on habitual control in patients, which may contribute to individual behavioral biases [Ersche et al., 2016, Lim et al., 2019, Heinz et al., 2019].

Based on these findings, we test whether habitual control can emerge through repeated action-sequence selection. Specifically, we ask whether participants can acquire a habitual response tendency in one context while maintaining goal-directed planning in another, and whether the BCC model can account for these behavioral dynamics. Moreover, we ask how participants adapt their behavior when switching between contexts that favor habitual control or require goal-directed planning. We investigate these questions using a combined computational and experimental approach with participant-specific model fitting.

We introduce a context-dependent sequential decision-making paradigm and apply the BCC model to test whether the observed behavioral dynamics can be captured in terms of habitual sequence priors, goal-directed evaluation, and context inference.

During acquisition, we expected repeated execution of the same optimal sequence to produce a performance advantage in the habit context, reflected in faster and more accurate responding. If this repetition establishes a persistent sequence tendency, changing the optimal sequence should initially impair performance in the habit context relative to the planning context. As participants adapt to the new contingencies, we expected this performance disadvantage to diminish and eventually give way to a renewed habit-context benefit. Because trials from the two contexts were interleaved, we further expected context switches to incur performance costs relative to context repetitions. Finally, we tested whether the BCC model could account for these behavioral dynamics in terms of learned sequence priors, goal-directed evaluation, and context inference.

## 2 Methods

### 2.1 Experimental Design

We now introduce the experiment, which is named *Two Galaxies Task (TGT)* and is an adaptation of a previously published forward planning task [Sass et al., 2025, Steffen et al., 2023]. Participants can adaptively switch between planning (goaldirected) and habitual (automated) responses in the form of action sequences to optimize the cost-benefit ratio in the respective context. The experimental design is based on the model’s assumption that habit formation is context-specific.

In the experiment, participants solve a three-step optimization problem on each trial. The task requires participants to navigate a spaceship through planetary systems with the goal of maximizing the amount of fuel collected as reward. Within a time limit of three seconds, participants have to enter a sequence of three decisions. The selected sequence determines the spaceship’s path through the planetary system. Screenshots of example trials are provided in Figure 5 in Appendix A.1. On each trial, one planet is randomly assigned as the starting position. The starting position itself does not affect the score. From this position, participants can choose between two possible actions: a flight to the nearest planet in the clockwise direction, or a flight to the second-nearest planet in the clockwise direction. We refer to these actions as a *short* flight and a *long* flight, respectively. The first action moves the spaceship to a new planet, from which the second action is again a short or a long flight to another planet. Each trial consists of a sequence of three such binary decisions, and the spaceship moves three times within each trial. Figure 1 depicts the state transition matrix in graph form. All transitions are deterministic.

**Figure 1:**
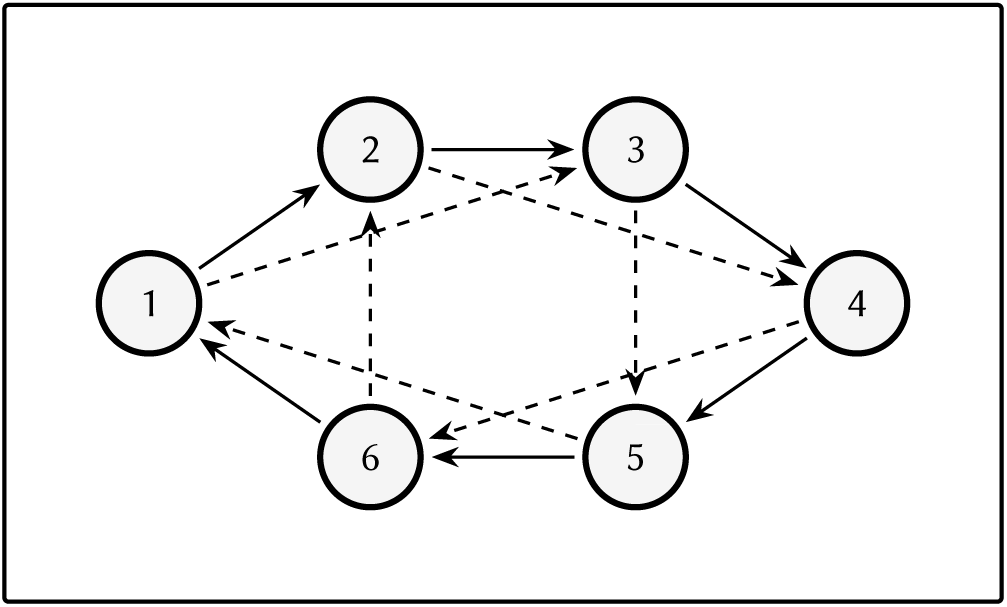
State transition matrix represented as a directed graph. Circles depict the states, i.e., the planet positions. Solid and dashed lines indicate short and long flight transitions, respectively.

The spaceship only starts moving through the planetary system after all three decisions have been entered. Consequently, participants have to plan the complete action sequence before observing the resulting path. Planning *on the fly* is therefore not possible. There are three green squares in the middle of the screen, as seen in Figure 5, that indicate the number of decisions that still have to be entered. In the example screenshots, all three decisions have yet to be selected by the participant, which is why there are still all three green squares visible. With each step entered, one of the squares disappears. Corrections are not possible.

There are three types of planets: blue, beige, and purple, associated with typical rewards of +1, 0, and −1 fuel points, respectively. Rewards are probabilistic, with each planet yielding its typical reward with a probability of 90%. In the remaining 10% of cases, blue and purple planets yield 0 fuel points instead of their typical reward. Beige planets occasionally yield a positive or negative reward instead of 0, with a probability of 5% each for +1 and −1 fuel points.

Since each of the three decisions can be either a short or a long flight, there are 2^3^ = 8 possible *action sequences*. A possible action sequence is for instance (1, 0, 1), where 0 encodes a short flight and 1 encodes a long flight. All trials are generated such that there is always one unique optimal action sequence. We quantify trial-wise performance using two outcome measures: the time required to enter the action sequence, and a binary *optimal choice* variable indicating whether the optimal decision sequence was selected.

Participants encounter two different contexts, implemented as different trial types. The current context is indicated by the background color of the trial. Importantly, participants were not informed that the background color was predictive of the underlying task structure.

When the background is green, one specific action sequence is always optimal. We refer to these trials as *habit trials*. When the background is brown, the optimal action sequence can be any of the eight possible decision sequences, with all sequences occurring equally often. We refer to these trials as *planning trials*. An example of a planning trial is shown in Figure 5B in Appendix A.1. The optimal decision sequence in this trial would be (0, 1, 0).

Thus, the two contexts differ in the structure of the optimal action sequence. In the habit context, the same action sequence is repeatedly optimal across trials. Repeated execution of this sequence is intended to induce a sequential habit, hence to induce a tendency for a specific automatic response. In the planning context, by contrast, participants have to flexibly plan ahead on each trial to identify the currently optimal action sequence.

The experiment is arranged into 10 *blocks* of 48 trials each, and every block consists of 24 *habit* and 24 *planning trials*. Trials from the two contexts are interleaved within each block by randomizing their order, with the restriction that the same context cannot occur more than four times consecutively. This design allows participants to repeatedly switch between a habit-like response mode and goal-directed planning while maintaining an equal number of trials from both contexts within each block. The participants were allowed to take breaks between each block. This block structure maintains the balance between contexts throughout the experiment.

The experiment is conducted over two consecutive days at approximately the same time of day (± 1 hour). On the first day, participants complete four blocks, and on the second day, they complete six blocks, as illustrated in Figure 2.

**Figure 2:**
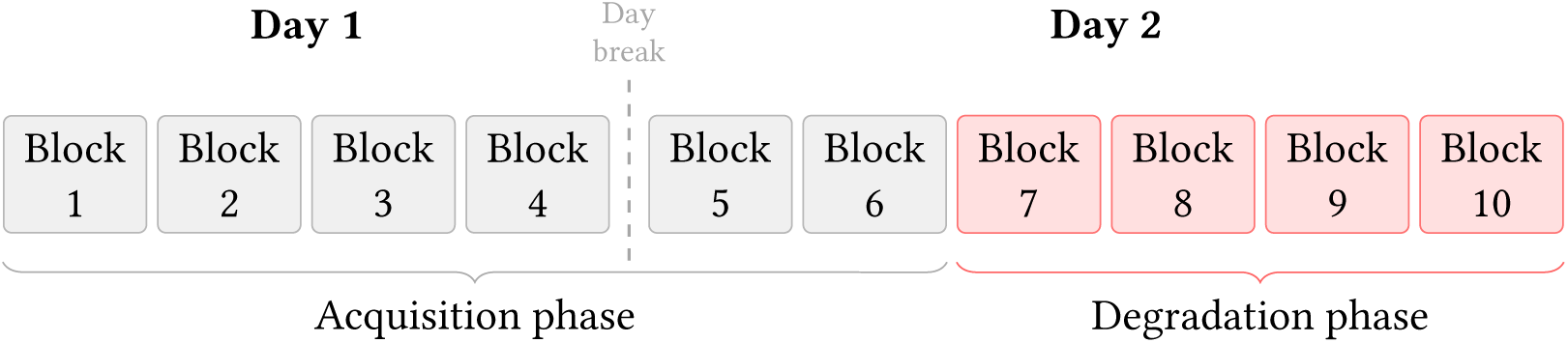
Block structure in experimental design.

The first six blocks during day 1 and day 2 constitute the *acquisition phase*. During this phase, the action sequence (0, 1, 1) is optimal on every trial in the habit context. As noted earlier, we expect that repeated execution of the same sequence in the habit context will lead to increasingly efficient and less deliberative responding. This is expected to result in a performance advantage for habit trials relative to planning trials, because the optimal sequence in the habit context can increasingly be retrieved or repeated rather than computed anew on each trial.

### 2.2 Hypotheses about Performance Measures

**Hypothesis 1.** During the acquisition phase, participants show better performance in the habit condition compared to the planning condition, reflected in faster reaction times and higher optimal-choice rates.

Since the hypothesis concerns both optimal-choice rates and reaction times, we fit separate mixed-effects models for the two outcome measures. The *block* was included as a centered continuous predictor, denoted by *block_c_*, in order to capture changes across the acquisition phase. Participant-specific baseline differences were modeled using random intercepts. We specify all mixed-effects models using standard R formula syntax, see Appendix A.5 for an explanation. For optimal choices, we fit the logistic mixed-effects model

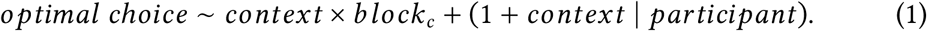

This model tests whether the probability of selecting the optimal action sequence differs between the habit and planning contexts, and whether this context effect changes across acquisition. Reaction times during the acquisition phase were analyzed using an analogous linear mixed-effects model with the same fixedand random-effects structure. Reaction times (RT) were log-transformed before analysis because the untransformed observations were positively skewed (skewness = 0.96). Log transformation substantially reduced this skewness (skewness = 0.20).

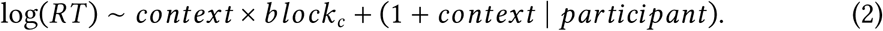

This model tests whether participants respond faster in the habit context than in the planning context, and whether this reaction-time difference changes across the acquisition phase.

The experiment includes a *degradation phase* in Blocks 7–10 on the second day. During this phase, the optimal action sequence in the habit context changes to (1, 1, 0), and the previously optimal action sequence is no longer optimal. If participants have acquired a sequential habit, they should be more likely to perseverate on the formerly optimal sequence despite the change in contingencies. The degradation phase therefore provides the condition for a behavioral test of whether repeated sequence execution in the habit context leads to reduced flexibility when the optimal sequence changes. Assuming that performance in the planning condition remains unaffected by degradation, we expect performance in the habit condition to initially fall below that in the planning condition.

**Hypothesis 2.** At the beginning of the degradation phase, participants show better performance in the planning condition compared to the habit condition, reflected in faster reaction times and higher optimal-choice rates.

Hypothesis 2 therefore concerns the relative performance difference between the habit and planning contexts at the onset of degradation. Because the same statistical analysis is used to test both this hypothesis and the following one, we describe the analysis after introducing the next hypothesis.

However, habits are not assumed to be entirely inflexible. Behavior may initially persist despite changes in contingencies but subsequently adapt to the new task structure. Therefore, across the degradation phase, participants may gradually learn the new optimal sequence in the habit context. Consequently, we expect the initial habitrelated performance cost to decrease over time and, by the end of the degradation phase, to turn back into a performance benefit in the habit context compared to the planning context.

**Hypothesis 3.** By the end of the degradation phase, participants regain a performance benefit in the habit condition compared to the planning condition, reflected in faster reaction times and higher optimal-choice rates.

Accordingly, the relevant test concerns the context-by-stage interaction rather than a within-context increase in performance alone. We test the Hypotheses 2 and 3 using separate mixed-effects models for optimal choices and reaction times. The analyses include the final acquisition block (Block 6) and all four degradation blocks (Blocks 7–10). Block is represented by a categorical variable referred to as *stage*. Block 6 represents the *end of acquisition*, Block 7 the *start of degradation*, Blocks 8 and 9 the intermediate degradation stages, and Block 10 the *end of degradation*. Modeling stage as a categorical predictor allows performance to differ between blocks without imposing a linear trend across the degradation phase. Block 7 (start of degradation) is used as the reference stage, with the planning context as the reference condition.

Although all five stages are included in the models, the hypothesis-focused estimated contrasts concern three theoretically relevant stages, which are the end of acquisition, the start of degradation, and the end of degradation. Habit-versus-planning contrasts at the start and end of degradation directly evaluate Hypotheses 2 and 3, respectively. The corresponding contrast at the end of acquisition characterizes the context difference immediately before degradation begins. The model-implied outcomes for Blocks 8 and 9 are additionally examined to describe the intermediate progression through degradation.

To test Hypotheses 2 and 3 with respect to optimal choices, we fit a logistic mixed-effects model predicting the trial-wise optimal-choice indicator from context, stage, and their interaction, with a by-participant random intercept:

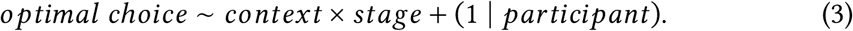

From this model, we estimate habit-versus-planning contrasts at the start and end of degradation. The contrast at the start tests for initial habit-context performance costs, whereas the contrast at the end tests whether a habit benefit has been regained.

In addition to these contrasts, we compare the end of acquisition with the start of degradation separately within each context. These contrasts evaluate whether optimal-choice rates change upon entering degradation in the planning and habit contexts. A direct difference-in-changes contrast then tests whether the transition-related change is greater in one context than in the other. This comparison is particularly relevant to Hypothesis 2, because it distinguishes a selective reduction in habit-context performance from a general change affecting both contexts. The planning-context contrast assesses whether there is evidence of a change in planning performance. To assess change across the degradation phase, we compare performances at the start and end of degradation separately within each context. A second difference-in-changes contrast tests whether this change differs between the planning and habit contexts. Blocks 8 and 9 are included in the categorical stage model and are reported descriptively as intermediate optimal-choice probabilities.

Reaction times are analyzed analogously using a linear mixed-effects model. The model predicts log-transformed reaction times from context, categorical stage, and their interaction, with a by-participant random intercept:

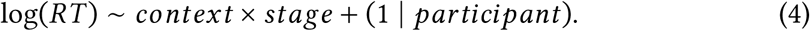

Habit-versus-planning contrasts are estimated at the end of acquisition and at the start and end of degradation. Within each context, additional contrasts quantify the change from the end of acquisition to the start of degradation and from the start to the end of degradation. Direct difference-in-changes contrasts test whether these changes differ between contexts. To complement the analysis of Hypothesis 2, we conducted a supplementary analysis of non-optimal choices during the first five habit trials of the degradation phase. We conjecture that the action sequence reinforced during acquisition briefly persists after degradation begins and is therefore overrepresented among erroneous choices relative to the other six non-optimal sequences.

Beyond differences in performance between contexts during the acquisition phase and respective changes induced by degradation, the TGT allows us to examine how performance is affected when the context changes from one trial to the next. In conventional task-switching experiments, alternating between tasks commonly produces switch costs, reflected in lower accuracy and longer reaction times on trials after a task switch [Rogers and Monsell, 1995, Ritz et al., 2026]. The TGT similarly involves transitions between the planning and habit contexts across successive trials. We therefore hypothesize that switching between these two contexts incurs performance costs, reflected in lower optimal-choice rates and longer reaction times.

To test this hypothesis, we classify each trial according to whether the context changes relative to the immediately preceding trial within the same block. Trials following a context switch are defined as a *switch trials*, whereas trials following the same context are defined as *repeat trials*. Because participants could take a break between blocks, we excluded the first trial of each block to avoid potential post-break effects.

**Hypothesis 4.** During the acquisition phase, switch trials are associated with fewer optimal choices and longer reaction times than repeat trials.

To test Hypothesis 4, we analyze trials from the acquisition phase using separate mixed-effects models for optimal choices and reaction time. The logistic mixed-effects model predicts trial-wise optimal choices from context, switch status, and their interaction. It includes a by-participant random intercept and a by-participant random slope for switch status:

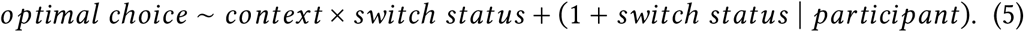

The corresponding linear mixed-effects model predicts log-transformed reaction times using the same fixed-and random-effects structure:

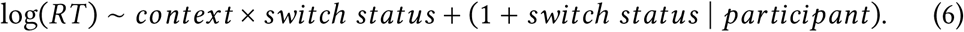

These models estimate the switch cost within each context, while the interaction tests whether the magnitude of the switch cost differs between the planning and habit contexts. We additionally estimate the switch cost averaged across both contexts.

### 2.3 Bayesian Modeling

Next, we outline the key principles of the BCC model underlying our Bayesian modeling approach. For further details, we refer to [Schwöbel et al., 2021]. We use the Bayesian inference framework at two levels. First, in the BCC, the agent performs Bayesian inference over contexts and action sequences. Second, we perform Bayesian inference over participant-specific model parameters when fitting the model to behavioral data. The model formalizes action selection as inference over latent contexts and candidate action sequences in a *hierarchical partially observable Markov decision process*. At each point in time *t*, the agent selects an action sequence and maintains beliefs about the hidden context *c_t_*. Both the environmental contingencies and the agent’s learned habits are context dependent.

An action sequence *π* is evaluated through two complementary components. First, the agent forms a context-specific prior over action sequences, *P*(*π* ∣ *c_t_*), which captures habitual control. This prior reflects the tendency to repeat action sequences that have been selected frequently in the same context in the past. Second, the agent evaluates the expected outcomes of an action sequence through a likelihood term, *P*(*r* ∣ *π*, *c_t_*), which captures goal-directed control by quantifying how well an action sequence is expected to lead to preferred outcomes *r*. action sequence selection is then sampled from the posterior distribution

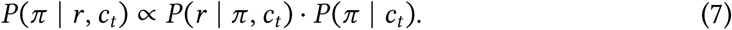

In this formulation, habitual and goal-directed control are not implemented as distinct systems with an additional arbitration mechanism. Instead, both influences are integrated directly through Bayesian inference. The prior over action sequences contributes the habitual bias, whereas the likelihood corresponds to the goal-directed evaluation of future outcomes for given action sequences and action selection natively balances the two by invoking Bayes’ theorem.

The current context is not directly observed, and therefore the agent must infer it from available data. Formally, this corresponds to maintaining a posterior over contexts, *P*(*c_t_* ∣ *d_t_*), where *d_t_* denotes the information available up to time *t*. The number of possible contexts was specified a priori as three. The posterior over action sequences is effectively weighted by uncertainty about the current context. More precisely, action sequence beliefs can be expressed as marginalization over contexts

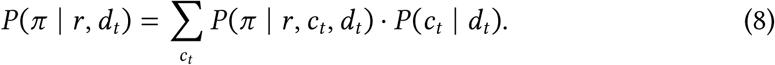

Consequently, a learned prior over action sequences only exerts a strong influence when the corresponding context is inferred to be active.

We consider three model parameters: a *goal weight δ*, a *habit weight* P, and a *context stability bias φ*. The habit weight parameter P controls how rapidly repeated action sequence selection is incorporated into the context-specific action sequence prior. In particular, we introduce weights for both the prior distribution *P*(*π* ∣ *c_t_*) and the likelihood *P*(*r* ∣ *π*, *c_t_*), where the latter resembles the typical inverse decision temperature, see e.g. [Daw et al., 2011, Friston et al., 2015]. Thus, we define

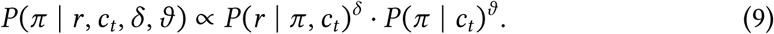

The goal weight *δ* controls the stochasticity of goal-directed action sequence selection, and the habit weight *θ* controls the stochasticity of the habitual prior. To see this, the posterior over action sequences can be written in terms of action sequence log-scores,

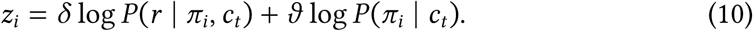

The final choice probabilities are then obtained by applying a softmax function:

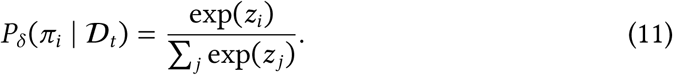

For numerical reasons, this parameter definition slightly differs from the one presented in [Schwöbel et al., 2021]. Lastly, the *context stability bias φ* determines how strongly the agent expects the latent context to persist across time. With three latent contexts, the corresponding self-transition probability is 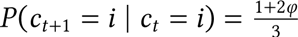. A strong bias toward context stability makes the agent less likely to infer that the task contingencies have changed, whereas a weaker stability bias allows the agent to switch more readily to a different inferred context when observations become surprising. This parameter is interesting because it can describe a persistent inference that the previously relevant context remains active, which might play a key role in addiction.

On the level of data analysis, we use Bayesian statistical inference to estimate participant-specific model parameters from observed behavioral data. That is, given a dataset *D_i_* from participant *i*, we infer a posterior over parameters *θ_i_*

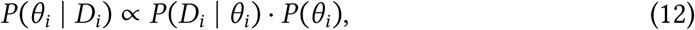

where *θ_i_* may include the parameters *δ*, P, and *φ*. To assess whether individual differences in context inference contribute to habit formation, we compared two variants of the BCC model. In both variants, *habit weight* P and *goal weight δ* were inferred as individual parameters. The reduced model used a fixed value for the *context stability bias φ*, whereas the extended model inferred this quantity as an additional individual parameter. In the first case, the value is fixed to the mean value of the estimates parameter values from the extended model.

In this model comparison, we generally test whether the BCC model can capture the transition from goal-directed to habitual behavior observed in the experimental data, and whether allowing for individual differences in the *context stability bias* improves the model fit. We refer to the model with only two free parameters, the goal weight *δ* and the habit weight P, as BCC-2P, and to the extended model variant that additionally includes the individual parameter *φ* as BCC-3P. We expect the additional parameter to improve model fit by capturing individual differences in the beliefs about context stability across trials.

**Hypothesis 5.** (a) The Bayesian contextual control model accounts for the choice behavior observed in the experimental data. (b) The BCC-3P model variant provides a better fit to the experimental data than the BCC-2P model variant.

The BCC model was implemented in python using *Torch* [Paszke et al., 2019], and the Bayesian inference of model parameters was implemented using *Pyro* [Bingham et al., 2019]. The code is openly available on GitHub [https://github.com/SSchwoebel/BalancingControl]. For the parameter inference, we used a version of the *PyBefit* toolbox [https://github.com/dimarkov/pybefit], which specifies hierarchical priors over parameters and calculates posterior distributions over model parameters using *stochastic variational inference (SVI)* [Hoffman et al., 2013] with *adaptive moment estimation (ADAM)* [Kingma and Ba, 2017] as optimization algorithm. To approximate warm-up experience, each reward prior was augmented with ten pseudocounts. We validated the model and its implementation with typical parameter and model recovery procedures as well as with posterior predictive checks, which we show in Appendix A.4. For parameter sweeps illustrating the effects of the three model parameters, see Figures 11 to 13 in Appendix A.4.

### 2.4 Technical Setup and Sample Description

The experiment was implemented in JavaScript and hosted on servers at Dresden University of Technology. Participants were recruited online via ORSEE [Greiner, 2015], which is the participant recruitment platform used by the Faculty of Psychology at Dresden University of Technology.

We collected data from 36 participants in an online experiment. Participants completed screening procedures for color vision impairments using the Ishihara test [Ishihara, 1917] and for motor impairments that could interfere with keyboard use. At the beginning of the experiment, participants completed a detailed instruction phase. During this phase, they were familiarized with the appearance of the game, its functionality, and the overall goal of the task. Several comprehension checks were included throughout the instructions to ensure that participants had understood the task correctly. Before the main experiment started, participants completed 20 warm-up trials on both days. All task instructions were presented in German.

Participants were assigned to one of two counterbalanced response-key mappings, with the key assignments for short and long flights reversed between the mappings Furthermore, to avoid order effects, we created four different trial-order configurations. These configurations were generated by shuffiing the complete set of blocks while preserving the structure of the degradation phase. Within each block, trials were shuffied as well, while keeping the constraint that no more than four consecutive trials belonged to the same context. After data collection, one participant was excluded because no responses were recorded in the last two blocks of the experiment. Therefore, the experiment resulted in a final sample of 35 participants. The distribution of trial-order configurations across participants as well as the keyboard mappings are reported in Table 6 in Appendix A.1. Participants ranged in age from 19 to 70 years (M = 29.37, SD = 12.31, median = 25), 26 participants were female (74%), and 33 participants held a higher education entrance qualification (94%).

At the end of the experiment, participants completed a debriefing questionnaire. They were asked whether they had noticed an association between the background colors and a the choice of a particular strategy. Interestingly, none of the 35 participants reported noticing the correct association.

## 3 Results

We first summarize task performance across blocks using reaction times and optimalchoice rates. We then report the mixed-effects analyses addressing Hypotheses 1–4, followed by the computational modeling results for the BCC model.

### 3.1 Task Performance

Figure 3 shows the mean reaction times and optimal-choice rates across blocks for the habit and planning contexts.

**Figure 3:**
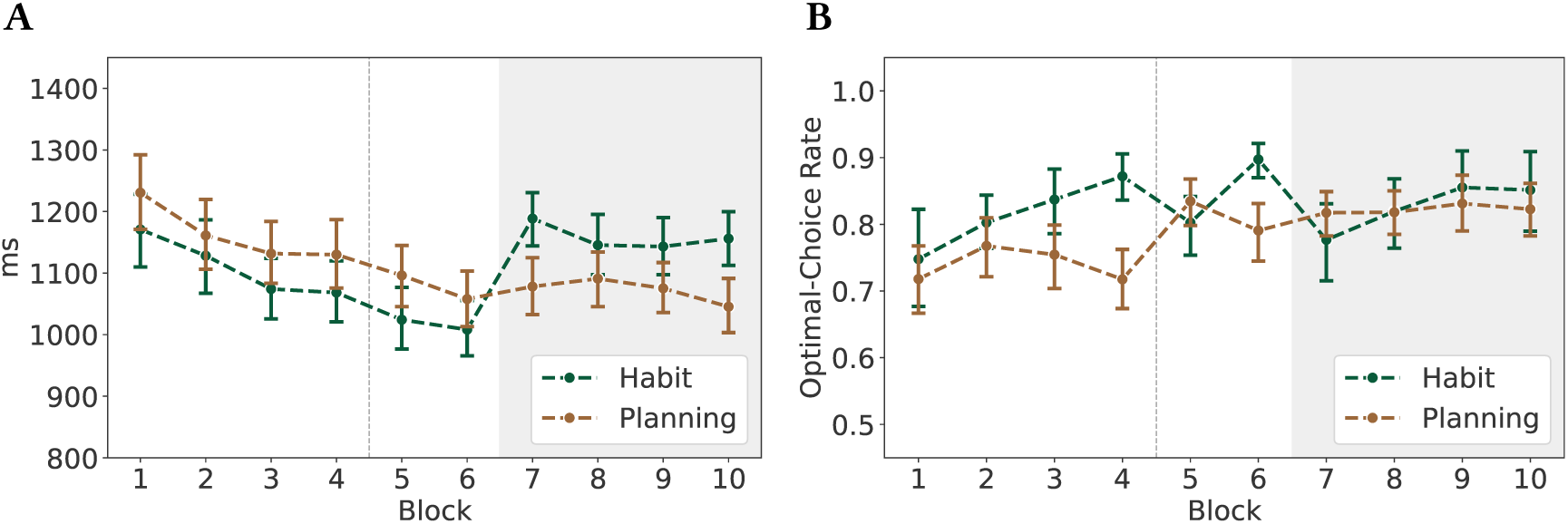
Task performance across blocks. **(A)** Mean reaction times across blocks for the habit and planning contexts. **(B)** Mean optimal-choice rates across blocks for the habit and planning contexts. In both panels, error bars indicate 95% confidence intervals. The dashed vertical line between Blocks 4 and 5 indicates the break between days, and the gray background indicates the degradation phase (Blocks 7–10). Lines connecting data points across blocks are included as visual aids only.

Since action sequences comprise three steps, there are three reaction times per trial. Figure 3A depicts the aggregated mean values of the first reaction time, and Figure 6A in Appendix A.2 shows the times for all three steps. The first reaction time is the informative measure and hereafter denoted as RT. Time-out rates resulting from the three-second response-time limit are shown in Figure 6B in Appendix A.2. Time-out trials were excluded from the calculation of aggregated performance measures and from all subsequent analyses. The optimal-choice rates, shown in Figure 3B, are the frequencies in which optimal sequences were chosen.

We refer to the performance advantage of the habit context over the planning context within each block as *habit benefit*. For the optimal-choice rates, the benefit was calculated as habit minus planning, and for reaction times, as planning minus habit, such that positive values consistently indicated better performance in the habit context. We refer to negative habit benefits as *habit costs*. Habit benefits for both performance measures are shown in Figure 7 in Appendix A.2. As a summarizing measure, Figure 8 in Appendix A.2 combines reaction times and optimal-choice rates using the *inverse efficiency score (IES)*. For each participant, block, and context, the IES was calculated by dividing the mean reaction time by the mean optimal choice rate. Therefore, lower values indicate more efficient performance.

#### Hypothesis 1: Habit Benefit During Acquisition

We first asked whether participants developed a performance advantage in the habit context during acquisition, as predicted by Hypothesis 1. Specifically, we examined whether optimal-choice rates were higher and reaction times shorter in the habit context than in the planning context.

**Optimal-Choice Rates** Participants were more likely to make an optimal response in the habit context than in the planning context (*β* = 0.480, *SE* = 0.061, *z* = 7.865, *p* < .001, *OR* = 1.616). This corresponds to approximately 62% higher odds of an optimal choice in the habit than in the planning context at the midpoint of the acquisition.

Furthermore, the optimal-choices increased across acquisition in both contexts, but the increase was stronger in the habit context. In the planning context, the odds of an optimal response increased significantly across acquisition blocks (*β* = 0.090, *SE* = 0.021, *z* = 4.378, *p* < .001, *OR* = 1.094). Each additional block was associated with an approximately 9.4% increase in the odds of an optimal response. This block-related change differed significantly between contexts, which is indicated by the positive context-by-block interaction (*β* = 0.068, *SE* = 0.032, *z* = 2.166, *p* = .030, *OR* = 1.071). Specifically, the habit-versus-planning odds ratio increased by approximately 7.1% with each additional block. Thus, the estimated habit benefit increased over the course of acquisition.

The complete estimates for the logistic mixed-effects model specified in Equation (1) are reported in Table 7 in Appendix A.3.

**Reaction Times** Reaction times were shorter in the habit context than in the planning context at the midpoint of the acquisition (*β* = −0.050, *SE* = 0.007, *t* = −7.224, *p* < .001.). The model-estimated reaction times were approximately 1041 ms in the habit context and 1095 ms in the planning context, which corresponds to a difference of 53 ms, or 4.9%.

Reaction times decreased across acquisition in the planning context (*β* = −0.025, *SE* = 0.002, *t* = −13.778, *p* < .001) by approximately 2.5% per block. The context-byblock interaction was not significant (*β* = −0.003, *SE* = 0.003, *t* = −1.097, *p* = .273). Thus, there was no evidence that the habit-versus-planning difference in reaction time changed over the acquisition blocks. Participants became faster over the acquisition phase and responded faster in the habit context than in the planning context, but the size of the estimated habit benefit did not clearly increase across blocks.

The complete estimates for the mixed-effects model for the log-transformed reaction times specified in Equation (2) are reported in Table 8 in Appendix A.3.

#### Hypotheses 2 and 3: Performance Decline and Recovery During Degradation

We next asked how performance changed when the previously reinforced action sequence ceased to be optimal. Hypothesis 2 predicted an initial habit cost, with lower performance in the habit than in the planning context at the onset of degradation, whereas Hypothesis 3 predicted that this disadvantage would diminish and a habit benefit would re-emerge by the end of degradation. We again examine optimal choice rates first and then turn to reaction times.

**Optimal-Choice Rates** Optimal-choice performance showed the expected initial habit cost at the onset of degradation. Participants were less likely to select the optimal action sequence in the habit context than in the planning context, which supports Hypothesis 2. Over the course of degradation, performance in the habit context recovered relative to the planning context. By the end of degradation, optimal-choice probability was numerically higher in the habit context, but this difference did not reach statistical significance, providing only limited support for Hypothesis 3.

The detailed model estimates for Equation (3) are reported in Table 9 in Appendix A.3. Table 1 summarizes the model-implied habit-versus-planning contrasts and estimated probabilities for each context at the end of acquisition, the start of degradation, and the end of degradation. The probabilities and contrasts for Blocks 8 and 9 can be found in Table 10 in Appendix A.3.

**Table 1:** Model-implied probabilities of an optimal choice across stages and estimated habit–planning contrasts. The Diff. (pp) column reports habit-minusplanning differences in percentage points. Contrast estimates are reported on the log-odds scale, with positive values and odds ratios above one indicating higher optimal-choice probabilities in the habit context.

| Stage | Planning | Habit | Diff. (pp) | Estimate | Odds ratio | *p* |
|---|---|---|---|---|---|---|
| End acquisition | .815 | .917 | +10.2 | 0.922 | 2.513 | < .001 |
| Start degradation | .842 | .802 | −4.0 | −0.276 | 0.759 | .033 |
| End degradation | .847 | .877 | +3.0 | 0.250 | 1.285 | .074 |

**Hypothesis 2: Initial Performance Decline** Supporting Hypothesis 2, the model-implied probability of an optimal choice was lower in the habit context (.802) than in the planning context (.842) at the start of degradation, which corresponds to a difference of −4.0 percentage points. This contrast was statistically significant (*β* = −0.276, *SE* = 0.129, *z* = −2.132, *p* = .033, *OR* = 0.759).

Furthermore, we examined whether the initial habit cost reflected a selective decline in habit-context performance rather than a general change across contexts. In the planning context, the probability of an optimal choice increased from .815 at the end of acquisition to .842 at the start of degradation, but this increase was not statistically significant (*β* = −0.189, *SE* = 0.130, *z* = −1.451, *p* = .147, *OR* = 0.828). Thus, there was no statistical evidence that planning-context performance changed upon entering the degradation phase. By contrast, the probability of an optimal choice in the habit context decreased from .917 at the end of acquisition to .802 at the start of degradation, and this decrease was statistically significant (*β* = −1.008, *SE* = 0.148, *z* = −6.795, *p* < .001, *OR* = 0.365). This corresponds to a reduction of approximately 11.5 percentage points.

Accordingly, the habit-versus-planning difference changed significantly from the end of acquisition to the start of degradation (*β* = 1.197, *SE* = 0.198, *z* = 6.061, *p* < .001). Together, these results indicate that the initial habit cost at the onset of the degradation was driven primarily by a pronounced decline in habit-context performance rather than by a statistically reliable change in planning-context performance.

A supplementary analysis examined whether the previous habit sequence was overrepresented among errors at the start of degradation (Table 11, Appendix A.3).

**Hypothesis 3: Recovery During Degradation** The data provide only limited support for Hypothesis 3, which predicted that the habit benefit in optimal-choice rates would recover by the end of the degradation phase. Although performance increased more strongly in the habit context than in the planning context during degradation, the habit benefit at the end of the phase was not statistically significant (*β* = 0.250, *SE* = 0.140, *z* = 1.788, *p* = .074, *OR* = 1.285). The model-implied probability of an optimal choice was numerically higher in the habit context (.877) than in the planning context (.847) by 3.0 percentage points.

We also examined whether performance in the habit context recovered relative to the planning context by the end of degradation. In the planning context, the probability of an optimal choice changed only slightly from .842 to .847. This change was not significant (*β* = 0.041, *SE* = 0.134, *z* = 0.307, *p* = .759, *OR* = 1.042). In the habit context, however, the probability of an optimal choice increased by 7.5 percentage points from .802 to .877, and this increase was statistically significant (*β* = 0.567, *SE* = 0.136, *z* = 4.184, *p* < .001, *OR* = 1.764).

Accordingly, the change from the start to the end of degradation differed significantly between contexts (*β* = 0.526, *SE* = 0.191, *z* = 2.760, *p* = .006). Thus, optimalchoice performance recovered in the habit context relative to the planning context. As shown in Table 10 in Appendix A.3, performance in the two contexts was already approximately equal by Block 8. Together, the results provide evidence for a performance improvement in the habit context relative to the planning context, but not for a statistically reliable recovery of a habit benefit by the end of degradation.

**Reaction Times** At the start of the degradation phase, participants responded more slowly in the habit context than in the planning context, which supports Hypothesis 2. This habit cost persisted at the end of the degradation, providing no support for the predicted recovery of the reaction-time benefit in Hypothesis 3.

The complete model estimates for Equation (4) are reported in Table 12 in Appendix A.3. Table 2 summarizes the model-implied geometric mean reaction times for each context, context differences in milliseconds, and habit-to-planning RT ratios at the end of acquisition, the start of degradation, and the end of degradation. The contrasts for Blocks 8 and 9 can be found in Table 13 in Appendix A.3.

**Table 2:**
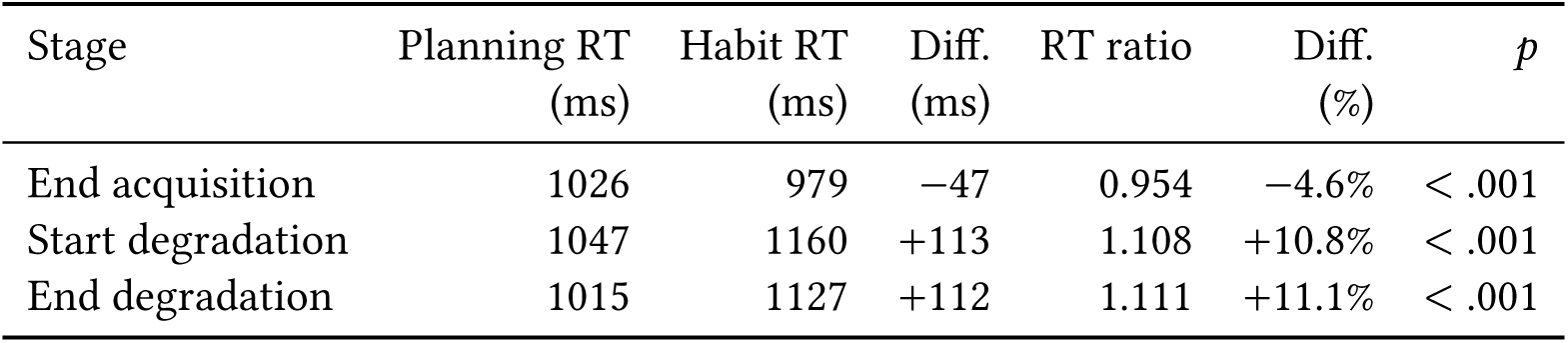
Model-based estimates of mean reaction times and habit-versus-planning differences across stages. Differences in milliseconds were calculated as habit minus planning. Positive values therefore indicate longer reaction times in the habit context. RT ratios are reported as habit-to-planning ratios, such that values above one indicate longer reaction times in the habit context.

| Stage | Planning RT<br>(ms) | Habit RT<br>(ms) | Diff.<br>(ms) | RT ratio | Diff.<br>(%) | *p* |
|---|---|---|---|---|---|---|
| End acquisition | 1026 | 979 | −47 | 0.954 | −4.6% | < .001 |
| Start degradation | 1047 | 1160 | +113 | 1.108 | +10.8% | < .001 |
| End degradation | 1015 | 1127 | +112 | 1.111 | +11.1% | < .001 |

**Hypothesis 2: Initial Performance Decline** The reaction-time results supported the initial performance decline predicted by Hypothesis 2. At the start of degradation, the model-implied geometric mean reaction time was longer in the habit context (1160 ms) than in the planning context (1047 ms), which corresponds to a difference of approximately 113 ms (10.8%). This contrast was statistically significant (*β* = 0.102, *SE* = 0.010, *t* = 10.157, *p* < .001, RT ratio = 1.108).

We next examined the transition from the end of acquisition to the start of degradation separately for each context. In the planning context, the estimated reaction time increased from 1026 ms at the end of acquisition to 1047 ms at the start of degradation. Reaction times in the planning context therefore increased by approximately 21 ms (2.1%) upon entering degradation. This contrast was statistically significant (*β* = 0.020, *SE* = 0.010, *t* = 2.016, *p* = .044, RT ratio = 1.021). In the habit context, the estimated reaction time increased from 979 ms at the end of acquisition to 1160 ms at the start of degradation. Thus, upon entering degradation, reaction times in the habit context increased by approximately 181 ms (18.5%), and this contrast was significant (*β* = 0.170, *SE* = 0.010, *t* = 16.858, *p* < .001, RT ratio = 1.185).

Accordingly, the reaction time increase from the end of acquisition to the start of degradation was significantly larger in the habit context than in the planning context (*β* = −0.150, *SE* = 0.014, *t* = −10.503, *p* < .001). The increase upon entering degradation was approximately 160 ms greater in the habit context than in the planning context. Together, the habit benefit at the end of acquisition reversed at the start of degradation, primarily driven by an increase in habit-context reaction times.

**Hypothesis 3: Recovery During Degradation** The reaction-time results provided no support for the recovery of the habit benefit predicted by Hypothesis 3. Although reaction times decreased significantly in the habit context over the course of degradation, they decreased to a similar proportional extent in the planning context. Consequently, reaction times remained significantly longer in the habit context (*β* = 0.105, *SE* = 0.010, *t* = 10.431, *p* < .001, RT ratio = 1.111). The model-based reaction-time estimates were approximately 1127 ms in the habit context and 1015 ms in the planning context, which is a difference of approximately 112 ms (11.1%).

We again examined the change from the start to the end of degradation. In the planning context, the estimated reaction time decreased from approximately 1047 ms to 1015 ms, which corresponds to a reduction of approximately 32 ms (3.1%). This decrease was statistically significant (*β* = −0.031, *SE* = 0.010, *t* = −3.089, *p* = .002, RT ratio = 0.969). In the habit context, the estimated reaction time decreased from approximately 1160 ms to 1127 ms, which is a reduction of approximately 33 ms (2.8%). This decrease was also statistically significant (*β* = −0.029, *SE* = 0.010, *t* = −2.831, *p* = .005, RT ratio = 0.972).

The proportional changes from the start to the end of degradation did not differ significantly between contexts (*β* = 0.003, *SE* = 0.014, *t* = 0.178, *p* = .859). Thus, there was no evidence of a stronger recovery in the habit context. The absolute reductions were also nearly identical, at approximately 33 ms in the habit context and 32 ms in the planning context.

#### Hypothesis 4: Context-Switching Costs

We next examined how performance was affected when the context changed from one trial to the next. Hypothesis 4 states that, during the acquisition phase, switch trials are associated with fewer optimal choices and longer reaction times than repeat trials. A *switch trial* was defined as a trial in which the context differed from that of the immediately preceding trial within the same block, whereas a *repeat trial* was preceded by a trial from the same context. We first present the results of the switchingcost analysis for optimal choices and then turn to reaction times.

**Optimal-Choice Rates** The optimal-choice probability was lower following context switches than following context repetitions, supporting Hypothesis 4. Table 3 summarizes the estimated optimal-choice probabilities and switch costs.

**Table 3:** Model-based estimates of optimal-choice probabilities and context-switching costs. Repeat and Switch denote estimated optimal-choice probabilities. Switching costs are expressed as probability differences (repeat minus switch) in per-centage points (pp) and as switch-to-repeat odds ratios (OR). Positive costs and odds ratios below one indicate reduced optimal choice following a context switch.

| Context | Repeat | Switch | Switch cost (pp) | OR | *p* |
|---|---|---|---|---|---|
| Planning | .809 | .754 | 5.45 [2.82, 8.08] | 0.73 | < .001 |
| Habit | .869 | .837 | 3.16 [0.95, 5.36] | 0.78 | .005 |

For planning trials, the estimated optimal-choice probability was lower following a habit trial (.754) than following a planning trial (.809), corresponding to a switch cost of 5.45 percentage points (*p* < .001). For habit trials, the estimated optimal-choice probability was likewise lower following a planning trial (.837) than following a habit trial (.869), corresponding to a switch cost of 3.16 percentage points (*p* = .005).

The estimated overall switch cost, with equal weighting of the two contexts, was 4.22 percentage points (*p* < .001). Although the estimated cost was numerically larger for planning than for habit trials, the context-by-transition interaction was not significant (*β* = 0.068, *SE* = 0.110, *z* = 0.614, *p* = .539). Thus, switching was associated with lower optimal-choice rates in both directions, without evidence that the size of this effect differed between the two contexts. The detailed estimates for the mixed-effects model specified in Equation (5) can be found in Table 14 in Appendix A.3.

As a supplementary analysis, we also examined whether planning errors following habit-to-planning switches reflected intrusions of the acquired habit action sequence. We analyzed 662 eligible error trials from all 35 participants. Of these errors, 139 consisted of the optimal habit sequence. A random-intercept logistic mixed-effects model estimated a probability of .208, which was significantly greater than the null probability of 1/7 = .143 (*z* = 4.351, one-sided *p* < .001).

**Reaction Times** The reaction-time results provided qualified support for Hypothesis 4, with a switch cost for transitions into the planning context but not for transitions into the habit context. Table 4 summarizes the estimated reaction times and corresponding context-switching costs.

**Table 4:**
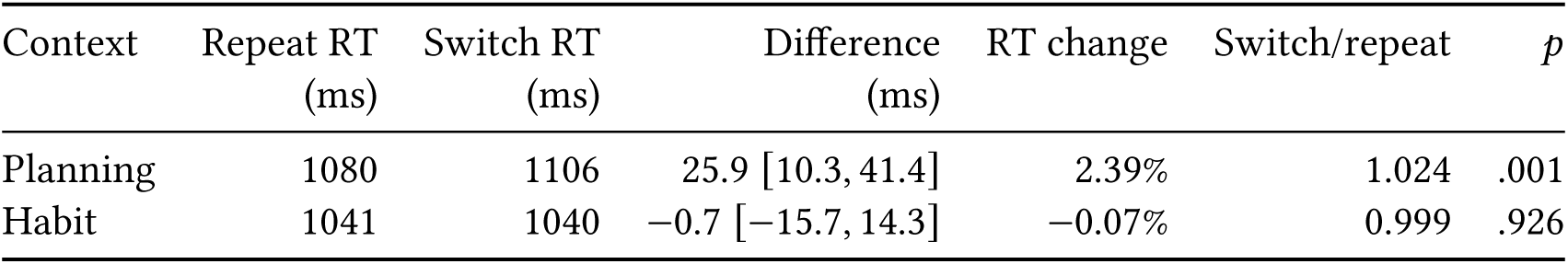
Model-based estimates of reaction times and context-switching costs. Repeat RT and Switch RT denote estimated reaction times by context. Switching costs are expressed as differences in milliseconds (switch minus repeat), percentage changes in reaction time, and switch-to-repeat ratios. Values in brackets are 95% confidence intervals for the differences. Positive differences and percentage changes, and ratios above one, indicate slower responses following a context switch.

For planning trials, the estimated mean reaction time was longer following a habit trial (1106 ms) than following a planning trial (1080 ms), which corresponds to a switch cost of 25.9 ms, or 2.39% (*p* = .001). For habit trials, the estimated mean reaction times were nearly identical following a planning trial (1040 ms) and following a habit trial (1041 ms), corresponding to a difference of −0.7 ms, or −0.07% (*p* = .926). Thus, there was no evidence of a reaction-time switch cost for habit trials.

The context-by-transition interaction was significant (*β* = −0.024, *SE* = 0.009, *t* = −2.629, *p* = .009), which indicates that the reaction-time switch effect was smaller in the habit than in the planning context. When the two contexts were weighted equally, the estimated overall switch cost was 12.3 ms (*p* = .037). The detailed estimates for the mixed-effects model specified in Equation (6) can be found in Table 15 in Appendix A.3.

### 3.2 Bayesian Modeling

We next examined whether allowing for individual differences in the context stability bias improved model fit. The BCC-2P model includes two free parameters, the habit weight P and the goal weight *δ*, whereas the BCC-3P model additionally includes the context stability bias parameter *φ*. In summary, both BCC model variants showed improved fit relative to the uniform random-choice null model, as indicated by mean pseudo-*R*^2^ values of approximately 0.57, consistent with Hypothesis 5(a). However, contrary to Hypothesis 5(b), allowing the context stability bias to vary across participants did not yield a clear improvement in model performance. Although WAIC numerically favored the BCC-3P model, the difference was not significant.

We first present the resulting parameter estimates before turning to the comparison of the two model variants. For the parameter inference, the context stability bias in the BCC-2P model was fixed at 0.46, which is the mean of the participant-level posterior mean estimates from the BCC-3P model. The same value was used for the BCC-2P model in the model and parameter recoveries and in the predictive posterior checks.

The negative ELBO stabilized across all empirical inferences and model and parameter recovery runs. Figure 4 summarizes the inferred individual model parameters. Model and parameter recovery analyses as well as the predictive posterior checks are reported in Appendix A.4. Correlations between the inferred model parameters are shown in Figure 15 in Appendix A.4.

**Figure 4:**
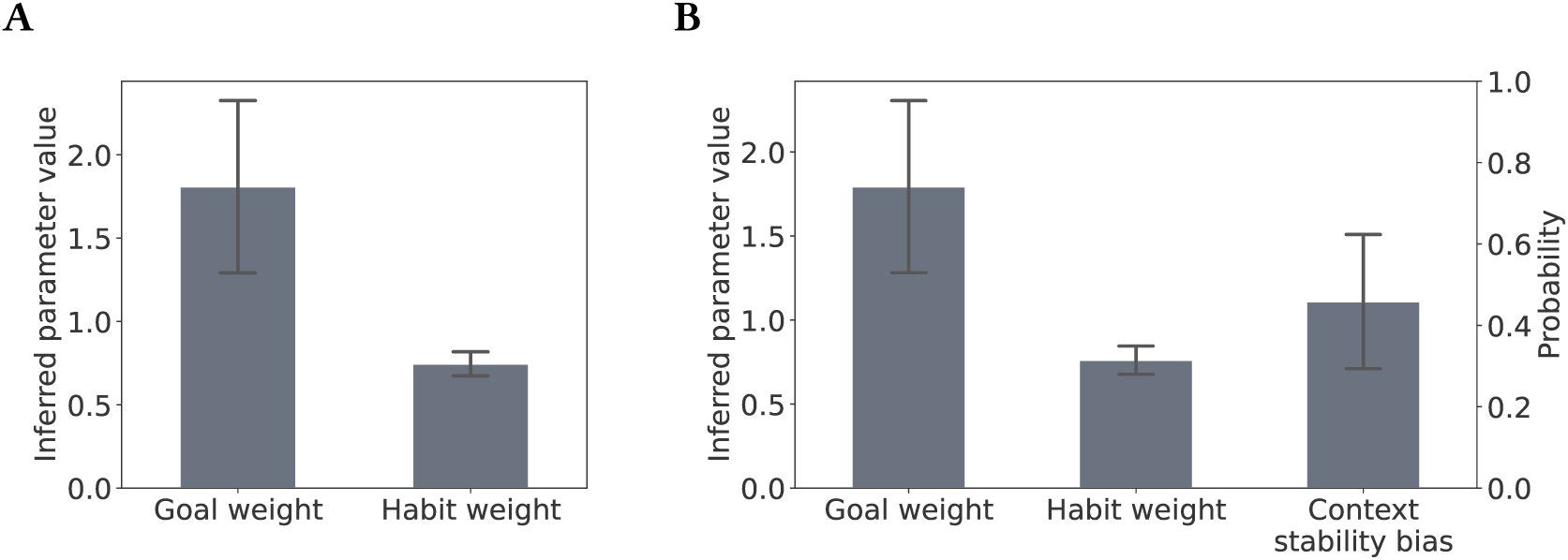
Inferred parameter values for the BCC-2P and BCC-3P model variants. **(A)** Inferred parameters for the BCC-2P model. **(B)** Inferred parameters for the BCC-3P model. Bars indicate the mean of participant-level posterior mean estimates, and error bars indicate the standard deviation across participants. The goal and habit weights are plotted against the left y-axis, and the context stability bias is plotted against the right y-axis.

We compared the two model variants using the widely applicable information criterion (WAIC), where lower values indicate better estimated out-of-sample predictive performance [Watanabe, 2010]. Invalid trials were excluded from both the fitted and null likelihoods and from the WAIC calculations. Thus, all scores were based on the same set of valid trials.

On average, the BCC-3P model had a slightly lower WAIC than the BCC-2P model, with mean values of 842.51 and 843.39, respectively. The mean WAIC difference, defined as BCC-3P minus BCC-2P, was −0.89 (95% CI [−1.79, 0.01]). This difference did not reach statistical significance in a paired-samples *t*-test (*t*(34) = −2.00, *p* = .053, Cohen’s *d_z_* = −0.34). Because participant-level scores shared information through the hierarchical fit, the test and confidence interval were interpreted as exploratory. Moreover, the BCC-3P model provided the better WAIC score for 22 of the 35 participants, whereas the BCC-2P model provided the better score for 13 participants.

An analysis of descriptive WAIC weights based on softmax(−WAIC/2) also showed a small numerical advantage for the BCC-3P model. The mean participantlevel weights were 0.418 for the BCC-2P model and 0.582 for the BCC-3P model.

Model fits were additionally evaluated using a pseudo-*R*^2^ index [McFadden, 1974], calculated as 1 − l_model_/l_null_. The null model corresponded to uniform random choice, with *p* = 0.5 at each valid choice point. The mean fitted log likelihoods were −415.26 for the BCC-2P model and −414.08 for the BCC-3P model, compared with a mean null log likelihood of −965.93. The BCC-2P model yielded a mean pseudo-*R*^2^ of 0.567 (*SD* = 0.134), and the BCC-3P model yielded a mean of 0.569 (*SD* = 0.135). Thus, adding the context stability bias parameter resulted in only a negligible improvement in likelihood-based fit relative to the null model. Table 5 summarizes the results of the comparison between the two model variants.

**Table 5:**
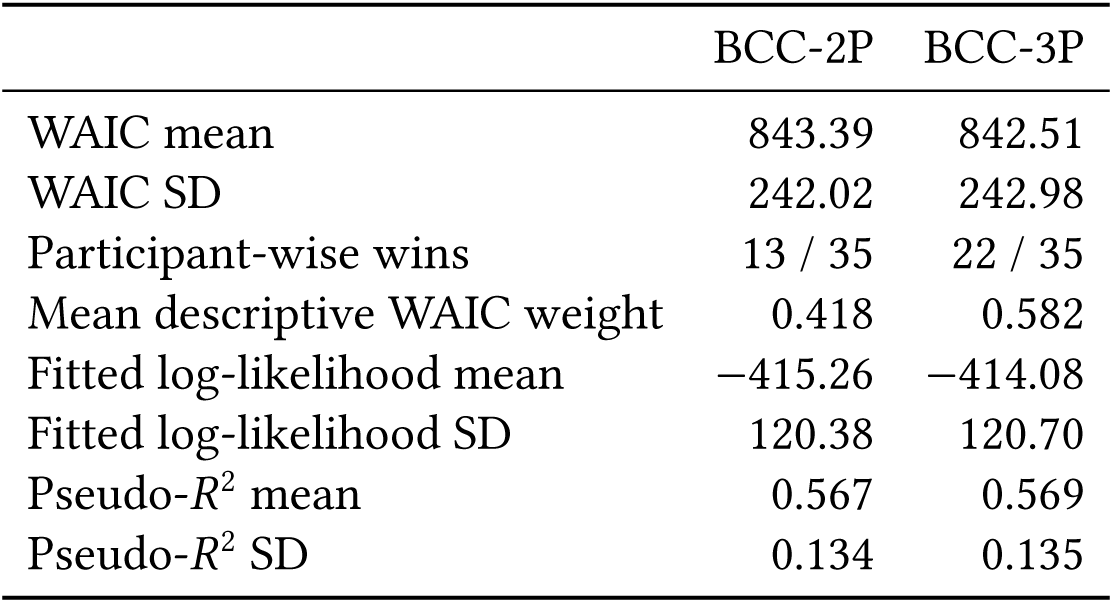
Model comparison between the BCC-2P and BCC-3P model variants. Lower WAIC values indicate better estimated predictive performance. Invalid trials were excluded from all likelihood and WAIC calculations. SD denotes the between-participant sample standard deviation.

|  | BCC-2P | BCC-3P |
|---|---|---|
| WAIC mean | 843.39 | 842.51 |
| WAIC SD | 242.02 | 242.98 |
| Participant-wise wins | 13 / 35 | 22 / 35 |
| Mean descriptive WAIC weight | 0.418 | 0.582 |
| Fitted log-likelihood mean | −415.26 | −414.08 |
| Fitted log-likelihood SD | 120.38 | 120.70 |
| Pseudo-*R*<sup>2</sup> mean | 0.567 | 0.569 |
| Pseudo-*R*<sup>2</sup> SD | 0.134 | 0.135 |

Overall, the model comparison provides no clear evidence that the additional context stability bias parameter improved model performance. Although WAIC and the descriptive weights numerically favored the BCC-3P model, the WAIC difference was small and the pseudo-*R*^2^ values were nearly identical. Parameter recovery for the context stability bias was also limited, with a correlation of approximately *r* = .40 between the true and inferred values, which suggests that individual differences in this parameter should be interpreted with caution.

Together, these findings support Hypothesis 5(a), with both BCC model variants fitting participant choices better than the uniform random-choice null model. Support for Hypothesis 5(b) was limited, with no clear improvement in model performance from allowing the context stability bias to vary across participants.

## 4 Discussion

The present study investigated the acquisition of context-dependent habits and adaptation to changed contingencies within the Two Galaxies Task paradigm. The task combines repeated selection of a stable optimal action sequence with an interleaved context in which optimal sequences change and planning is required. This design allows us to examine sequence repetition, contextual control, and behavioral flexibility within the same experimental setting.

To provide a mechanistic account of participant choices, we applied the Bayesian contextual control model, which describes action selection through Bayesian inference within the active inference framework [Schwöbel et al., 2021]. The model combines learned priors over action sequences with goal-directed evaluation, conditional on beliefs about the current context. The BCC model provides a computational framework for linking the observed behavioral patterns to the joint influence of learned action-sequence priors, goal-directed evaluation, and context inference. This makes it possible to examine repetition-based habitual tendencies within a unified account of context-dependent sequential choice.

### 4.1 Behavioral Findings

As hypothesized, we found that participants responded faster and more accurately in the habit condition than in the planning condition. This performance benefit indicates automatization when repeating a stable sequence. These findings are consistent with repetition-based accounts of habit formation [Miller et al., 2019] and with proposals that familiar sequences support efficient responding through hierarchical organization or chunking [Dezfouli and Balleine, 2012, Dezfouli et al., 2014]. Related experimental work has shown that repeated sequence execution improves performance while biasing subsequent responses toward the learned sequence [Frölich et al., 2023]. The present task examines these benefits while interleaving planning trials in which the optimal action sequence changes across trials and must therefore be determined anew. However, faster and more accurate responding during acquisition could also reflect motor fluency or goal-directed retrieval of a familiar solution. The subsequent change in contingencies therefore provides a more informative test of whether the previously practiced sequence continues to bias behavior when it is no longer optimal.

At the beginning of the degradation phase, participants responded more slowly and made fewer optimal choices in the habit condition than in the planning condition, supporting *Hypothesis 2*. In the TGT, *degradation* denotes a change in task contingencies that makes a different sequence optimal, and it does not denote classical contingency degradation, which weakens the relation between an action and its outcome. The decline in performance after the contingency change is consistent with the previously learned sequence interfering with selection of the newly optimal sequence [Miller et al., 2019, Frölich et al., 2023]. Taken together, the performance advantage during acquisition and the subsequent cost after the contingency change suggest a trade-off: repeated execution of the same sequence facilitates performance while contingencies remain stable, but can impede adaptation once those contingencies change. Evidence for persistent execution of the previously optimal sequence was more limited. A significant increase in errors attributable to that sequence was detected only on the first habit trial of degradation. The subsequent performance cost may therefore reflect additional effort to suppress the habitual response or evaluate alternatives, rather than continued overt selection of the old sequence. The behavioral data do not directly distinguish these processes, and an initial adaptation cost alone does not imply the outcome insensitivity that is central to classical definitions of habit [Dickinson, 1985, Balleine and Dickinson, 1998].

The results provided limited support for *Hypothesis 3*, which predicted a renewed habit-context benefit by the end of degradation. Optimal-choice performance recovered relative to the planning condition, but a significant advantage was not regained, and reaction times even showed no corresponding recovery. This pattern is consistent with participants learning to select the newly optimal sequence while continuing to evaluate alternatives or resolve competition from the trained sequence. It is compatible with accounts in which goal-directed selection and habitual tendencies coexist [Dezfouli et al., 2014, Frölich et al., 2023]. However, longer reaction times could also reflect caution or uncertainty following the change. Recovery in optimal choices alone therefore establishes neither the extinction of the original tendency nor the acquisition of a replacement habit.

The context-switching results provide qualified support for *Hypothesis 4*. Switching reduced optimal-choice probabilities in both directions. Reaction times increased when participants switched from the habit context to the planning context, whereas no detectable cost occurred in the opposite direction. This asymmetric reactiontime pattern suggests that transition demands might be lower when switching from the planning to the habit context. Task-switching research similarly links performance costs to preparation demands and interference from preceding task states [Rogers and Monsell, 1995, Ritz et al., 2026].

One possible explanation for this asymmetry is that entering the planning context required renewed evaluation of alternative sequences, whereas entering the habit context allowed relatively rapid retrieval of a habitual sequence. This interpretation is consistent with the proposed role of context in activating learned response tendencies [Wood and Rünger, 2016]. It also aligns with a resource rationality perspective, according to which relying on learned sequences can reduce computational demands when the expected benefits of further planning do not justify its costs [Kool et al., 2017]. However, despite the absence of a reaction-time cost when switching into the habit context, optimal-choice probability was reduced following switches in both directions. Thus, retrieval of the familiar sequence may have reduced the time cost of switching into the habit context without eliminating switching-related errors.

None of the 35 participants reported noticing an association between background color and the execution of a particular sequence. The effects therefore occurred without explicit awareness of this association. Taken together, the repetition benefits, adaptation costs, and switching effects motivate a computational account in which acquired sequence tendencies interact with contextual information.

These behavioral findings should be viewed in light of how the TGT differs from established paradigms. The standard two-step task distinguishes model-based and model-free contributions over two decision stages, but does not include a dedicated period of stable sequence acquisition followed by a separate test of adaptation, or explicitly alternate contexts favoring habitual responding and continued planning [Daw et al., 2011]. The difficulty of reliably inducing outcome-insensitive habits in humans further motivates complementary approaches to studying habit formation [de Wit et al., 2018]. Another distinction is that participants in the TGT enter the complete action sequence before its execution, whereas the two-step task and the actionsequence task provide feedback after each action [Daw et al., 2011, Frölich et al., 2023].

### 4.2 Computational Modeling

The computational results support that the BCC model can account for the observed choices (*Hypothesis 5(a)*). Within this framework, habitual tendencies are represented by learned priors over action sequences and combined with goal-directed evaluation through Bayesian inference [Schwöbel et al., 2021]. Repetition strengthens the prior associated with a sequence in a given context. Stable contingencies therefore allow a consistent sequence prior to develop, whereas changing optimal sequences require continued sensitivity to expected outcomes. The model offers an account of how these influences can contribute jointly to choices in the habit and planning contexts.

This account differs from influential reinforcement learning approaches that associate habitual behavior with model-free learning of cached action values [Daw et al., 2005, Dolan and Dayan, 2013]. The habitual component of BCC represents a repetition-based preference for previously executed sequences, rather than a cached estimate of their reward value. It shares this assumption with value-free accounts of habit learning [Miller et al., 2019]. Its representation of sequences also connects it to hierarchical accounts of action control [Dezfouli and Balleine, 2012, Dezfouli et al., 2014]. However, this conceptual similarity should not be taken as evidence that participants represented or executed the learned sequences as indivisible action chunks. The model also provides a distinct account of the interaction between habitual and goal-directed influences. Arbitration models allocate control according to the uncertainty or reliability of predictions from model-based and model-free systems [Daw et al., 2005, Lee et al., 2014]. In BCC, the interaction follows from combining habitual priors and goal-directed evaluation within the same inference procedure [Schwöbel et al., 2021]. Beliefs about the active context determine which priors and contingencies contribute to choice. This provides a computational interpretation of context-dependent habit expression [Wood and Rünger, 2016]. The present application extends the model’s theoretical account to participant-specific fitting, while its explanatory specificity remains to be tested against competing models.

*Hypothesis 5(b)*, which predicted better performance of the BCC-3P model than the BCC-2P model, was not supported. Estimating a participant-specific context selftransition probability did not improve model performance sufficiently to favor the additional parameter. One possible explanation is that background color provided an informative context cue, reducing the contribution of individual differences in prior beliefs about context stability. The parameter sweep is consistent with this interpretation, but does not entail that participants relied primarily on color. Both model variants include context inference, and the comparison indicates that its additional participant-specific flexibility was not warranted by these data.

The BCC framework also distinguishes possible sources of behavioral persistence. Continued selection of a previously optimal sequence may reflect a strong habitual prior, a weaker influence of goal-directed evaluation, or, when context representations favor different sequences, continued belief in the previously active context. Whether these potential sources of behavioral persistence can be distinguished empirically, however, depends on how reliably the corresponding model parameters can be recovered. Parameter recovery was very good for goal weight and habit weight, supporting the estimation of these two parameters under the tested conditions. Recovery of context transition bias was moderate, which suggests a greater need for caution when interpreting individual differences in beliefs about context stability.

Lastly, the fitted parameters showed greater between-participant variability in goal weight than in habit weight. This pattern raises the possibility that individual differences in the present task were expressed more strongly through goal-directed evaluation than through the weighting of habitual priors. Although both parameters showed very good recovery, their observed variability is not necessarily directly comparable, because it also depends on estimation uncertainty. The finding should therefore remain exploratory rather than be interpreted as evidence that goal-directed control is intrinsically more variable across individuals than habitual control.

### 4.3 Limitations and Future Directions

The main interpretive limitation is that the present design cannot determine whether the observed repetition-related sequence tendencies constitute habits in the classical sense of outcome-insensitive control. Acquisition benefits and adaptation costs provide evidence of repetition-related changes in performance, but they can also arise from motor learning or retrieval of familiar solutions.

As noted above, the degradation phase changes which sequence is optimal, rather than selectively weakening an action–outcome contingency. A complementary outcome-devaluation or classical contingency-degradation test would help determine whether the acquired tendencies meet established criteria for habitual control [Dickinson, 1985, Balleine and Dickinson, 1998]. This distinction is especially relevant given the difficulty of demonstrating training-induced outcome insensitivity in humans [de Wit et al., 2018].

The experiment also covered a relatively short time scale and may primarily capture early habit formation. Longer acquisition periods would test whether repetition produces more persistent sequence tendencies. Longer degradation periods would help determine whether an adaptation eventually restores a performance benefit in both optimal-choice rates and reaction times.

A further limitation is that the present design does not allow us to isolate the contribution of context inference from the effects of repeated sequence execution. Because the context cue and the underlying sequence structure were systematically linked, future studies should manipulate context cues independently of sequence contingencies to test if context information modulates the expression of learned action tendencies.

Finally, another direction for future work is to examine these mechanisms in clinical populations with SUD. Habit-based accounts of addiction motivate testing whether clinical groups differ in sequence persistence, adaptation, or context switching [Everitt and Robbins, 2005, Everitt and Robbins, 2016, Ersche et al., 2016]. However, impaired flexibility can reflect reduced goal-directed control as well as stronger habitual tendencies, and recent work on alcohol use emphasizes the need for careful translation of these constructs [de Wit et al., 2018, Giannone et al., 2024]. The BCC framework offers hypotheses about these alternatives, including altered context inference, whose evaluation requires reliable behavioral measures and identifiable model parameters. To extend this work to the clinical domain, we are currently conducting a study using this paradigm in a clinical sample.

### 4.4 Summary

The present findings are consistent with the acquisition of context-dependent habits through repeated selection of stable action sequences. Repetition improved performance during acquisition and was followed by an initial cost when contingencies changed. The BCC model accounted for participant choices through the interaction of sequence priors, goal-directed evaluation, and context inference. Adding a participantspecific context stability parameter did not improve model performance relative to the more parsimonious BCC-2P model. Together, the task and modeling approach provide a basis for studying repetition and adaptation in context-dependent sequential decision making. Comparisons with competing models and complementary tests of habitual control will help to establish the specificity of this account.

## Funding Acknowledgments

This research was funded by the German Research Foundation (Deutsche Forschungs-gemeinschaft, DFG project numbers 390696704 (EXC 2050/2), 402170461 (TRR 265), 454245598 (IRTG 2773), and 521379614 (TRR 393)).

## Data, Code, and Material Availability

The model, analysis scripts, and experiment code are available at https://gitlab.com/willi.hieke/context-dependent-sequential-decision-making. The data supporting the findings of this study are available from the corresponding author upon reasonable request.

## Author Contributions

WH, SH, SS, AP, and MNS designed the experiment. SS developed the underlying model and provided the model code. WH and MNS formulated the hypotheses. WH, SH, and AP conducted the pilot studies. WH collected data from the final sample, adapted and applied the model code, and performed the model comparison. WH, AP, and MNS planned the statistical analyses of the behavioral outcomes, which WH performed. WH and MNS interpreted the results. MNS and SJK supervised the research and provided critical feedback on manuscript drafts. WH took the lead in writing the manuscript. All other authors reviewed the manuscript and approved the final version.

## Ethics Approval and Informed Consent

The study was approved by the ethics committee of the Dresden University of Technology (EK-570122022) and performed in accordance with relevant guidelines and regulations. All participants provided written informed consent before participation.

### Appendix

#### A.1 Experiment

**Figure 5:**
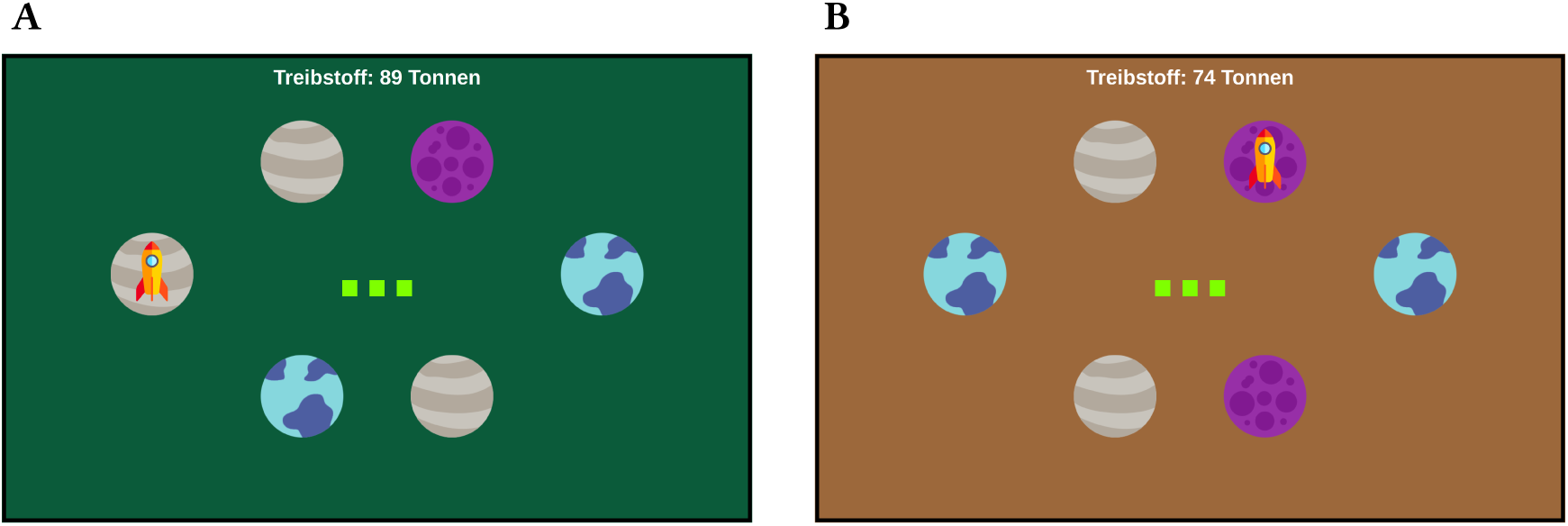
Example trials in the Two Galaxies Task. **(A)** An example habit trial. **(B)** An example planning trial.

**Table 6:** Distributions of trial configurations and key bindings.

| Trial configuration | Count | Key binding | Count |
|---|---|---|---|
| No. 1 | 10 | No. 1 | 20 |
| No. 2 | 11 | No. 2 | 15 |
| No. 3 | 7 |  |  |
| No. 4 | 7 |  |  |

#### A.2 Behavioral Readouts

**Figure 6:**
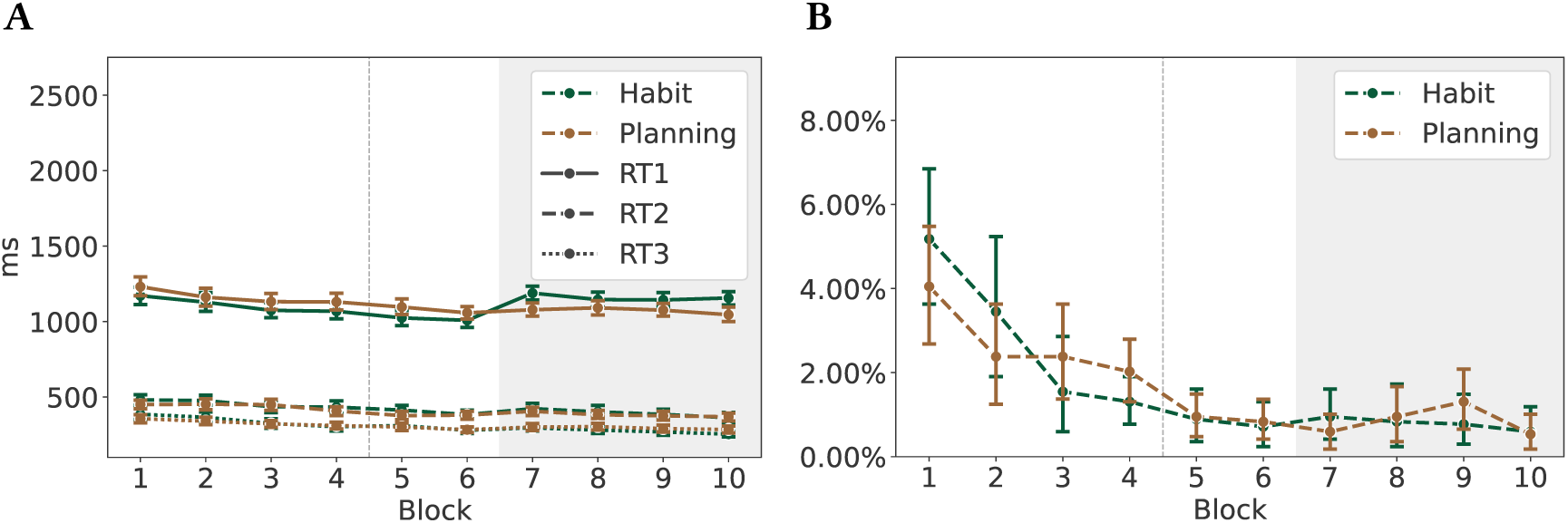
Reaction times 1–3 and time-out rates. **(A)** Reaction times 1–3 across blocks. **(B)** Time-out rates across blocks. The layout follows that of Figure 3.

**Figure 7:**
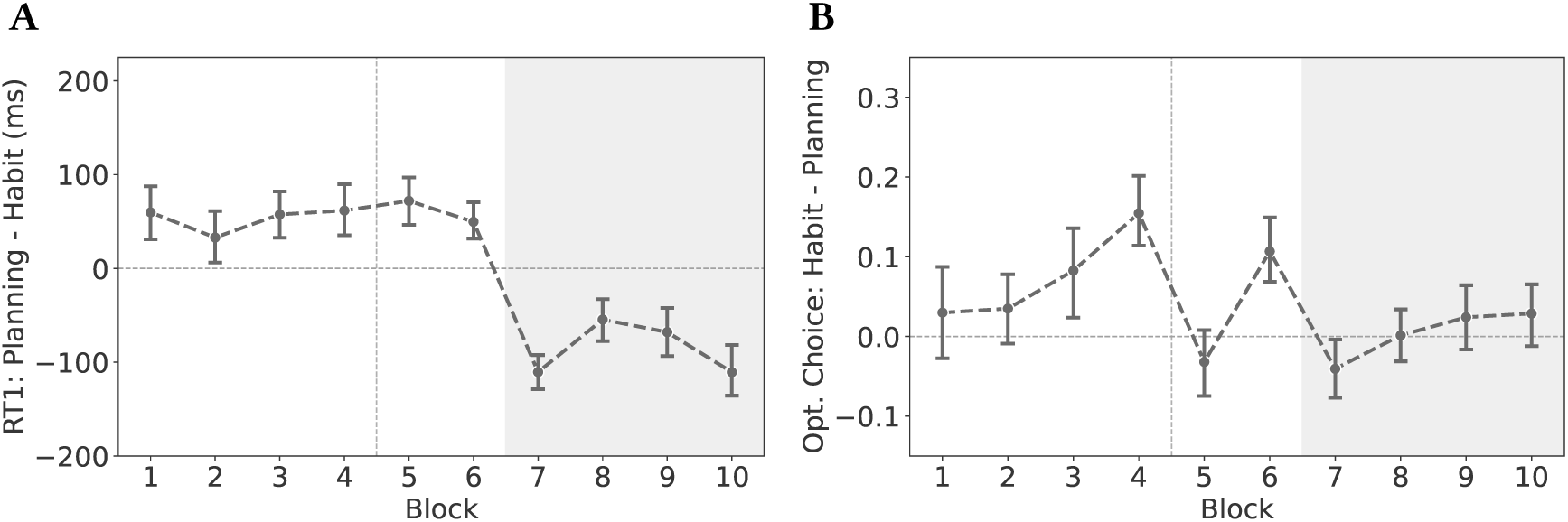
Habit benefits in reaction times and optimal-choice rates. **(A)** Mean reaction-time habit benefit across blocks. **(B)** Mean optimal-choice-rate habit benefit across blocks. The habit benefit in each block represents the mean of the context differences calculated separately for each participant. Positive reaction-time values indicate faster responses in the habit context than in the planning context, whereas positive optimal-choice-rate values indicate a higher optimal-choice rate in the habit context. The layout follows that of Figure 3.

**Figure 8:**
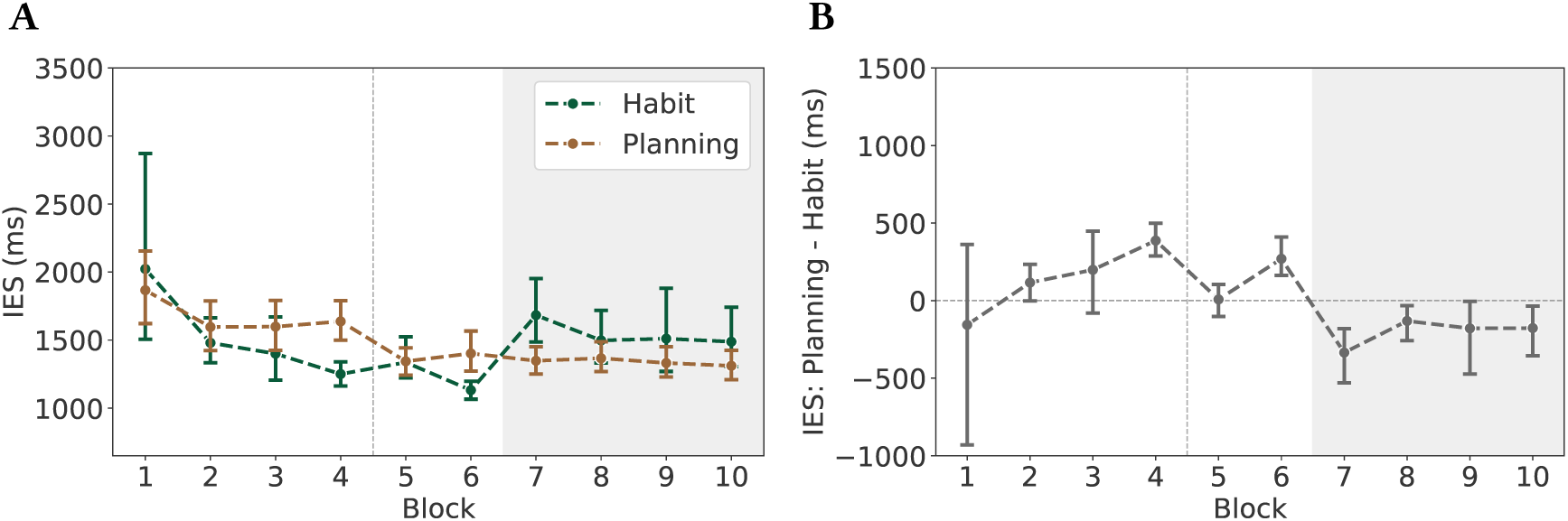
Inverse efficiency scores and corresponding habit benefits. **(A)** Mean IES across blocks for the habit and planning contexts. **(B)** Mean habit benefit in IES across blocks. For each participant and block, the IES was calculated separately for each context by dividing the mean reaction time by the mean optimal-choice rate. The resulting context differences were then averaged across participants. The layout follows that of Figure 3.

#### A.3 Mixed-Effects Models

**Table 7:** Logistic mixed-effects model predicting optimal choices during acquisition. The model is specified in Equation (1). Fixed-effect estimates are reported on the log-odds scale. The planning context serves as the reference category, and block was centered at its mean across acquisition trials. The exponentiated intercept therefore represents the predicted odds of an optimal choice in the planning context at the centered block, and the exponentiated context coefficient represents the habit-toplanning odds ratio at the centered block. The exponentiated block coefficient represents the odds ratio per one-block increase in the planning context. The exponentiated interaction represents the factor by which the habit-to-planning odds ratio changes per one-block increase.

| **Fixed effects** |  |  |  |  |  |
|---|---|---|---|---|---|
| Predictor | Estimate | SE | *z* | *p* | exp(*β*) |
| Intercept | 1.259 | 0.101 | 12.421 | < .001 | 3.521 |
| Context: habit | 0.480 | 0.061 | 7.865 | < .001 | 1.616 |
| Block, centered | 0.090 | 0.021 | 4.378 | < .001 | 1.094 |
| Context × Block | 0.068 | 0.032 | 2.166 | .030 | 1.071 |
| **Random effects** |  |  |  |  |  |
| Grouping factor | Effect | Variance | SD | Corr. |  |
| Participant | Intercept | 0.313 | 0.560 | – |  |
| Participant | Context: habit | 0.020 | 0.140 | .64 |  |
| **Model information** |  |  |  |  |  |
| Observations |  |  |  |  | 9,648 |
| Participants |  |  |  |  | 35 |
| Blocks |  |  |  |  | 1–6 |
| Marginal *R*<sup>2</sup> |  |  |  |  | .028 |
| Conditional *R*<sup>2</sup> |  |  |  |  | .127 |

**Table 8:** Linear mixed-effects model predicting log-transformed reaction times during acquisition. The model is specified in Equation (2). Fixed-effect estimates are reported on the log-RT scale. The planning context serves as the reference category, and block was centered at its mean across acquisition trials. The exponentiated intercept therefore represents the predicted geometric-mean reaction time in milliseconds in the planning context at the centered block, The exponentiated context coefficient represents the habit-to-planning RT ratio at the centered block. The exponentiated block coefficient represents the multiplicative change in RT per one-block increase in the planning context. The exponentiated interaction represents the factor by which the habit-to-planning RT ratio changes per one-block increase.

| **Fixed effects** |  |  |  |  |  |
|---|---|---|---|---|---|
| Predictor | Estimate | SE | *t* | *p* | exp(*β*) |
| Intercept | 6.998 | 0.022 | 314.556 | < .001 | 1094.477 |
| Context: habit | −0.050 | 0.007 | −7.224 | < .001 | 0.951 |
| Block, centered | −0.025 | 0.002 | −13.778 | < .001 | 0.975 |
| Context × Block | −0.003 | 0.003 | −1.097 | .273 | 0.997 |
| **Random effects** |  |  |  |  |  |
| Grouping factor | Effect | Variance | SD | Corr. |  |
| Participant | Intercept | 0.017 | 0.130 | – |  |
| Participant | Context: habit | 0.001 | 0.031 | .12 |  |
| Residual | Residual | 0.048 | 0.218 | – |  |
| **Model information** |  |  |  |  |  |
| Observations |  |  |  |  | 9,648 |
| Participants |  |  |  |  | 35 |
| Blocks |  |  |  |  | 1–6 |
| Marginal *R*<sup>2</sup> |  |  |  |  | .040 |
| Conditional *R*<sup>2</sup> |  |  |  |  | .303 |

**Table 9:** Logistic mixed-effects model predicting optimal choices at the end of acquisition and during degradation. The model is specified in Equation (3). Fixedeffect estimates are reported on the log-odds scale. The planning context and the start of degradation serve as the reference categories for context and stage, respectively. The exponentiated intercept therefore represents the predicted odds of an optimal choice in the planning context at the start of degradation, and the exponentiated context coefficient represents the habit-to-planning odds ratio at the start of degradation. The exponentiated stage coefficients represent the odds ratios for each stage relative to the start of degradation in the planning context. The exponentiated interaction coefficients represent the factors by which the habit-to-planning odds ratio differs at each stage relative to the start of degradation.

| **Fixed effects** |  |  |  |  |  |
| Predictor | Estimate | SE | *z* | *p* | exp(*β*) |
|---|---|---|---|---|---|
| Intercept | 1.673 | 0.160 | 10.424 | < .001 | 5.328 |
| Context: habit | −0.276 | 0.129 | −2.132 | .033 | 0.759 |
| Stage: end acquisition | −0.189 | 0.130 | −1.451 | .147 | 0.828 |
| Stage: degradation Block 8 | 0.005 | 0.134 | 0.037 | .970 | 1.005 |
| Stage: degradation Block 9 | 0.106 | 0.136 | 0.780 | .435 | 1.112 |
| Stage: end degradation | 0.041 | 0.134 | 0.307 | .759 | 1.042 |
| Context × Stage: end acquisition | 1.197 | 0.198 | 6.061 | < .001 | 3.312 |
| Context × Stage: degradation Block 8 | 0.300 | 0.187 | 1.604 | .109 | 1.349 |
| Context × Stage: degradation Block 9 | 0.481 | 0.193 | 2.495 | .013 | 1.617 |
| Context × Stage: end degradation | 0.526 | 0.191 | 2.760 | .006 | 1.692 |
| **Random effects** |  |  |  |  |  |
| Grouping factor | Effect | Variance | SD |  |  |
| Participant | Intercept | 0.584 | 0.764 |  |  |
| **Model information** |  |  |  |  |  |
| Observations |  |  |  |  | 8,264 |
| Participants |  |  |  |  | 35 |
| Blocks |  |  |  |  | 6−10 |
| Marginal *R*<sup>2</sup> |  |  |  |  | .019 |
| Conditional *R*<sup>2</sup> |  |  |  |  | .167 |

**Table 10:** Model-implied probabilities of an optimal choice and estimated habit-versus-planning contrasts at the end of acquisition and across all degradation blocks. The Diff. (pp) column reports differences between the habit and planning contexts in percentage points, calculated as habit minus planning. Contrast estimates are reported on the log-odds scale. Positive differences and estimates, as well as odds ratios above one, indicate higher optimal-choice probabilities in the habit context. Dashes indicate that no inferential context contrast was evaluated for that block. The intermediate-block estimates illustrate how performance developed during degradation.

| Stage | Planning | Habit | Diff. (pp) | Estimate | Odds ratio | *p* |
|---|---|---|---|---|---|---|
| End of acquisition | .815 | .917 | +10.2 | 0.922 | 2.513 | < .001 |
| Start of degradation | .842 | .802 | −4.0 | −0.276 | 0.759 | .033 |
| Degradation Block 8 | .843 | .846 | +0.3 | – | – | – |
| Degradation Block 9 | .856 | .879 | +2.3 | – | – | – |
| End of degradation | .847 | .877 | +3.0 | 0.250 | 1.285 | .074 |

##### Errors at Degradation Entry

As a supplementary analysis of the drop in optimalchoice rates at the start of the degradation phase, we examined whether the previously reinforced habit sequence (0, 1, 1) was disproportionately selected among erroneous choices in Block 7. Under the assumption that each of the seven non-optimal action sequences was equally likely, the old habit was expected to account for 1/7 = 14.29% of errors. The analysis was repeated for cumulative windows of the first one to five habit trials. Time-out trials were used for the definition of the windows but were excluded from the statistical analysis.

As shown in Table 11, evidence for perseveration of the old habit was strongest on the first habit trial. Of the 17 errors committed on this trial, 6 involved the old habit, which corresponds to 35.3%. This proportion was significantly greater than the chance level of 14.29% (*z* = 2.34, one-sided *p* = .010). When the window was extended to the first two habit trials, the old habit accounted for 6 of 23 errors (26.1%). Although this remained numerically above chance, the one-sided test did not reach the conventional significance threshold (*z* = 1.58, *p* = .057). Across the first three habit trials, the old habit accounted for 7 of 34 errors (20.6%), which was not significantly greater than chance (*z* = 1.04, *p* = .149). By the cumulative five-trial window, the old-habit error rate had declined to 14.0%, approximately matching the expected rate of 14.29%.

**Table 11:** Use of the formerly reinforced habit action sequence among erroneous choices during the beginning of degradation. The 95% confidence intervals and one-sided tests were obtained from intercept-only logistic models. The null probability was 1/7 = 14.29%. Because the analysis windows were cumulative, they were not statistically independent.

| Early window | Valid trials | Errors | Habit errors | Old-habit rate | *z* | *p* |
|---|---|---|---|---|---|---|
| First trial | 34 | 17 | 6 | 35.3% [16.8%, 59.6%] | 2.34 | .010 |
| First two trials | 67 | 23 | 6 | 26.1% [12.2%, 47.2%] | 1.58 | .057 |
| First three trials | 102 | 34 | 7 | 20.6% [10.1%, 37.3%] | 1.04 | .149 |
| First four trials | 136 | 39 | 7 | 17.9% [8.8%, 33.1%] | 0.65 | .257 |
| First five trials | 171 | 50 | 7 | 14.0% [6.8%, 26.6%] | −0.058 | .523 |

Overall, the results suggest that perseveration of the formerly reinforced habit was concentrated primarily on the first habit trial following degradation. As additional trials were included, the observed old-habit error rate declined and no longer differed significantly from chance.

**Table 12:** Linear mixed-effects model predicting log-transformed reaction times at the end of acquisition and during degradation. The model is specified in Equation (4). Fixed-effect estimates are reported on the log-RT scale. The planning context and the start of degradation serve as the reference categories for context and stage, respectively. The exponentiated intercept therefore represents the predicted geometric-mean reaction time in milliseconds in the planning context at the start of degradation. The exponentiated context coefficient represents the habit-to-planning RT ratio at the start of degradation. The exponentiated stage coefficients represent the RT ratios for each stage relative to the start of degradation in the planning context. The exponentiated interaction coefficients represent the factors by which the habitto-planning RT ratio differs at each stage relative to the start of degradation.

| **Fixed effects** |  |  |  |  |  |
|---|---|---|---|---|---|
| Predictor | Estimate | SE | *t* | *p* | exp(*β*) |
| Intercept | 6.954 | 0.021 | 335.240 | < .001 | 1046.971 |
| Context: habit | 0.102 | 0.010 | 10.157 | < .001 | 1.108 |
| Stage: end acquisition | −0.020 | 0.010 | −2.016 | .044 | 0.980 |
| Stage: degradation Block 8 | 0.011 | 0.010 | 1.069 | .285 | 1.011 |
| Stage: degradation Block 9 | −0.004 | 0.010 | −0.372 | .710 | 0.996 |
| Stage: end degradation | −0.031 | 0.010 | −3.089 | .002 | 0.969 |
| Context × Stage: end acquisition | −0.150 | 0.014 | −10.503 | < .001 | 0.861 |
| Context × Stage: degradation Block 8 | −0.049 | 0.014 | −3.444 | < .001 | 0.952 |
| Context × Stage: degradation Block 9 | −0.036 | 0.014 | −2.544 | .011 | 0.964 |
| Context × Stage: end degradation | 0.003 | 0.014 | 0.178 | .859 | 1.003 |
| **Random effects** |  |  |  |  |  |
| Grouping factor | Effect | Variance | SD |  |  |
| Participant | Intercept | 0.013 | 0.115 |  |  |
| Residual | Residual | 0.042 | 0.205 |  |  |
| **Model information** |  |  |  |  |  |
| Observations |  |  |  |  | 8,264 |
| Participants |  |  |  |  | 35 |
| Blocks |  |  |  |  | 6−10 |
| Marginal *R*<sup>2</sup> |  |  |  |  | .046 |
| Conditional *R*<sup>2</sup> |  |  |  |  | .275 |

**Table 13:** Model-implied geometric-mean reaction times and estimated habitversus-planning contrasts at the end of acquisition and across all degradation blocks. Differences in milliseconds are calculated as habit minus planning. Contrast estimates are reported on the log-RT scale. RT ratios are calculated as habit values divided by planning values. Positive differences and estimates, as well as RT ratios above one, indicate longer reaction times in the habit context. Dashes indicate that no inferential context contrast was evaluated for that block. The intermediate-block estimates illustrate how reaction times developed during degradation.

| Stage | Planning RT (ms) | Habit RT (ms) | Diff. (ms) | RT ratio | Diff. (%) | *p* |
|---|---|---|---|---|---|---|
| End of acquisition | 1026 | 979 | −47 | 0.954 | −4.6% | < .001 |
| Start of degradation | 1047 | 1160 | +113 | 1.108 | +10.8% | < .001 |
| Degradation Block 8 | 1058 | 1116 | +58 | 1.055 | +5.5% | – |
| Degradation Block 9 | 1043 | 1114 | +71 | 1.068 | +6.8% | – |
| End of degradation | 1015 | 1127 | +112 | 1.111 | +11.1% | < .001 |

**Table 14:** Logistic mixed-effects model predicting optimal choices as a function of context and transition status. The model is specified in Equation (5). Fixedeffect estimates are reported on the log-odds scale. The planning context and context repetition serve as the reference categories for context and transition status, respectively. The exponentiated intercept therefore represents the predicted odds of an optimal choice on repeat trials in the planning context. The exponentiated context coefficient represents the habit-to-planning odds ratio on repeat trials. The exponentiated transition coefficient represents the switch-to-repeat odds ratio in the planning context. The exponentiated interaction represents the factor by which the habit-to-planning odds ratio differs on switch trials relative to repeat trials.

| **Fixed effects** |  |  |  |  |  |
|---|---|---|---|---|---|
| Predictor | Estimate | SE | *z* | *p* | exp(*β*) |
| Intercept | 1.441 | 0.115 | 12.588 | < .001 | 4.226 |
| Context: habit | 0.450 | 0.085 | 5.292 | < .001 | 1.569 |
| Transition: switch | −0.321 | 0.076 | −4.206 | < .001 | 0.726 |
| Context × Transition | 0.068 | 0.110 | 0.614 | .539 | 1.070 |
| **Random effects** |  |  |  |  |  |
| Grouping factor | Effect | Variance | SD | Corr. |  |
| Participant | Intercept | 0.345 | 0.587 | – |  |
| Participant | Transition: switch | 0.013 | 0.116 | .22 |  |
| **Model information** |  |  |  |  |  |
| Observations |  |  |  |  | 9,465 |
| Participants |  |  |  |  | 35 |
| Blocks |  |  |  |  | 1–6 |

**Table 15:** Linear mixed-effects model predicting log-transformed reaction times as a function of context and transition status. The model is specified in Equation (6). Fixed-effect estimates are reported on the log-RT scale. The planning context and context repetition serve as the reference categories for context and transition status, respectively. The exponentiated intercept therefore represents the predicted geometric-mean reaction time in milliseconds on repeat trials in the planning context. The exponentiated context coefficient represents the habit-to-planning RT ratio on repeat trials. The exponentiated transition coefficient represents the switch-to-repeat RT ratio in the planning context. The exponentiated interaction represents the factor by which the habit-to-planning RT ratio differs on switch trials relative to repeat trials.

| **Fixed effects** |  |  |  |  |  |
|---|---|---|---|---|---|
| Predictor | Estimate | SE | *t* | *p* | exp(*β*) |
| Intercept | 6.985 | 0.023 | 304.312 | < .001 | 1079.849 |
| Context: habit | −0.037 | 0.007 | −5.310 | < .001 | 0.964 |
| Transition: switch | 0.024 | 0.007 | 3.381 | .001 | 1.024 |
| Context × Transition | −0.024 | 0.009 | −2.629 | .009 | 0.976 |
| **Random effects** |  |  |  |  |  |
| Grouping factor | Effect | Variance | SD | Corr. |  |
| Participant | Intercept | 0.01762 | 0.1327 | – |  |
| Participant | Transition: switch | 0.00024 | 0.0154 | −.11 |  |
| Residual | Residual | 0.04982 | 0.2232 | – |  |
| **Model information** |  |  |  |  |  |
| Observations |  |  |  |  | 9,465 |
| Participants |  |  |  |  | 35 |
| Blocks |  |  |  |  | 1–6 |

#### A.4 Bayesian Contextual Control Model

**Figure 9:**
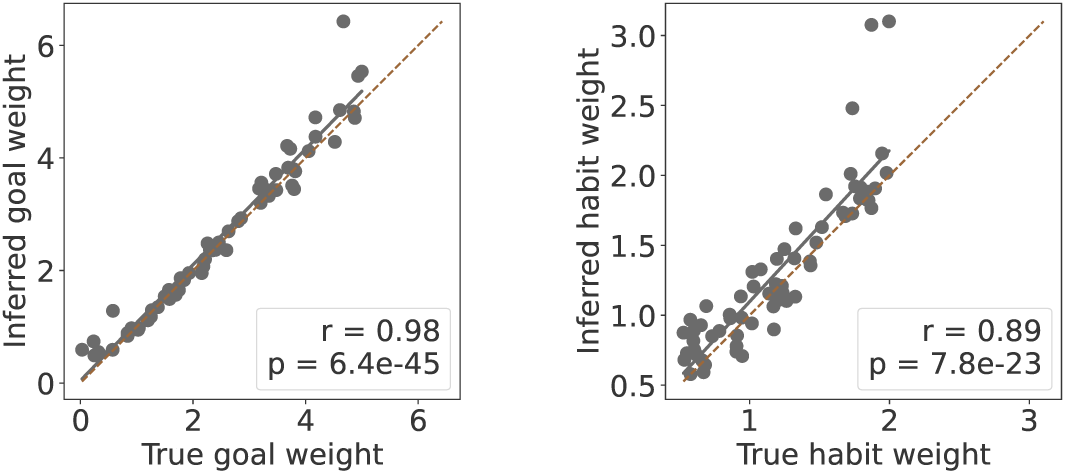
Parameter recovery for BCC-2P (n=64)

**Figure 10:**
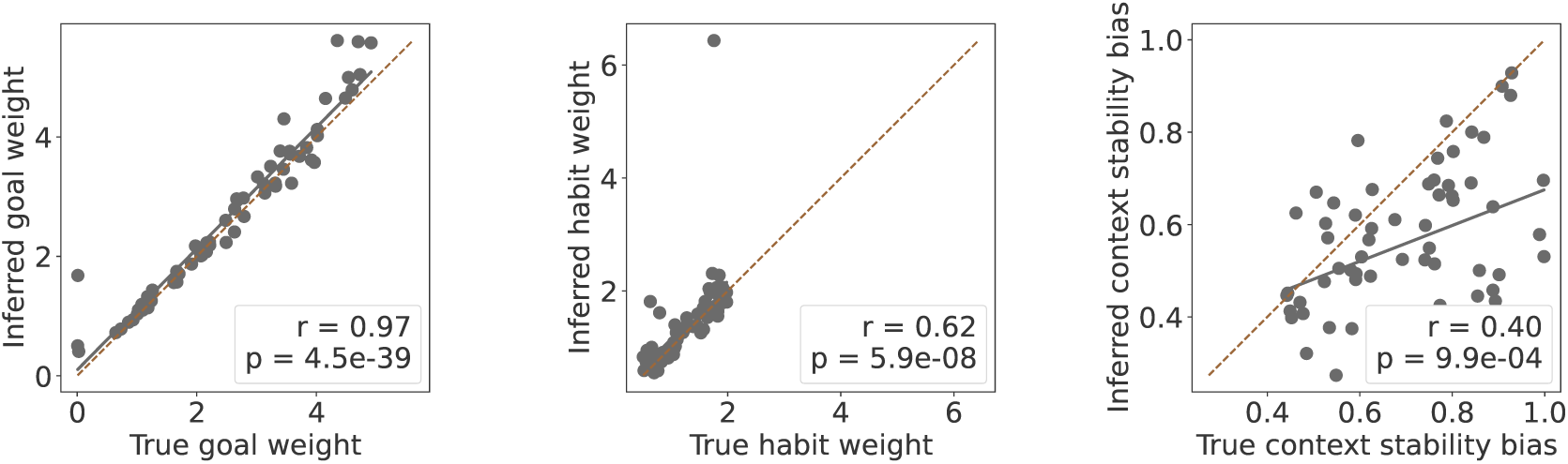
Parameter recovery for BCC-3P (n=64).

**Table 16:** Model recovery results for the BCC-2P and BCC-3P models. For each generative model, 10 datasets comprising 32 simulated agents each were generated. Rows indicate the generative model and columns indicate the recovered model based on the lower mean WAIC. Values show the proportion of recovery repetitions assigned to each model, with correct recoveries shown on the diagonal.

|  | Recovered model |  |
|---|---|---|
| Generative model | BCC-2P | BCC-3P |
| BCC-2P | **80%** | 20% |
| BCC-3P | 10% | **90%** |

**Figure 11:**
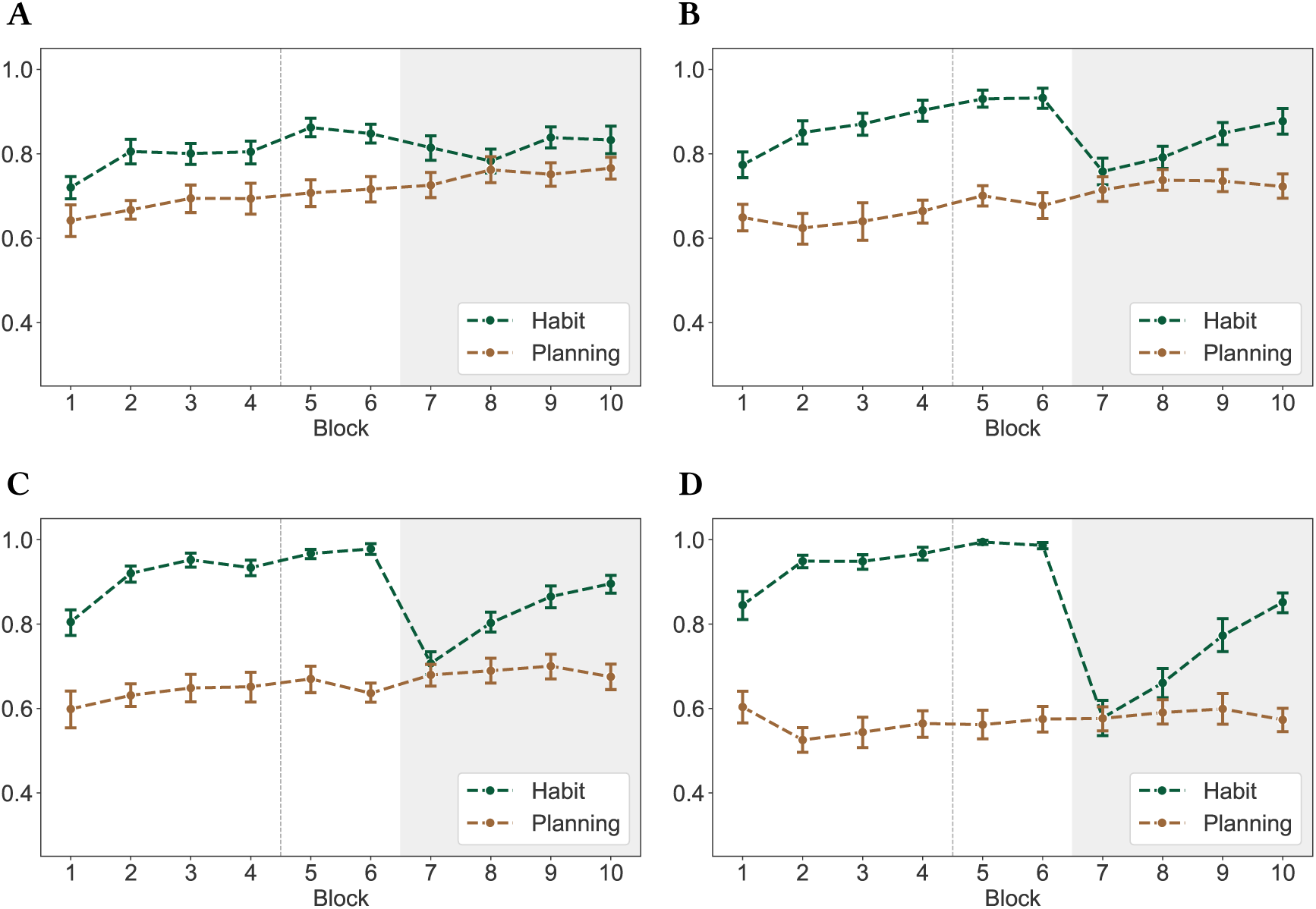
Optimal-choice rates across habit-weight values. **(A)** Simulation with P = 0.25. **(B)** Simulation with P = 0.75. **(C)** Simulation with P = 1.25. **(D)** Simulation with P = 1.75. Each parameter setting was simulated using 32 agents. The *goal weight δ* was fixed at 2, and the *context stability bias φ* was fixed at 0.5. The layout follows that of Figure 3.

**Figure 12:**
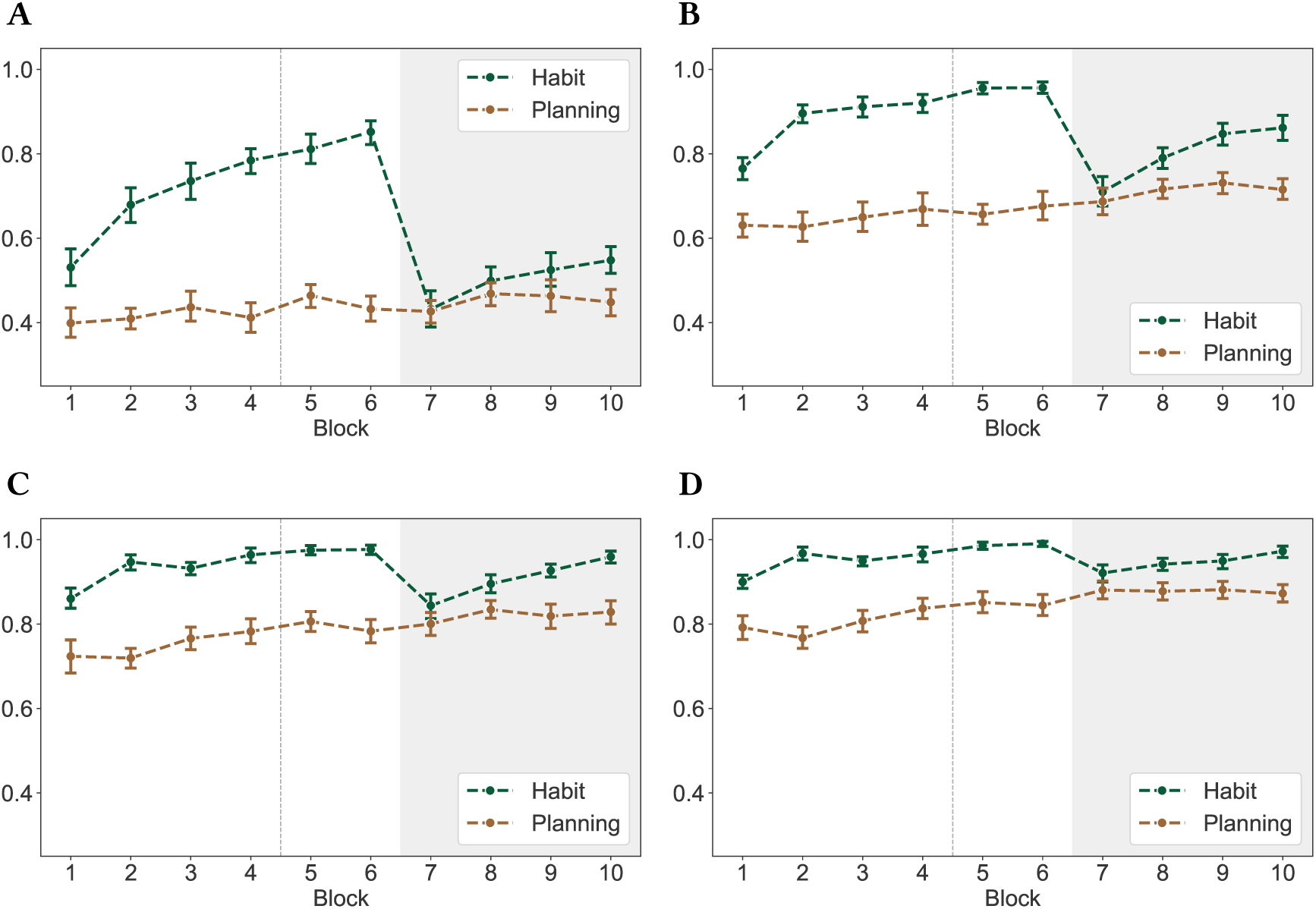
Optimal-choice rates across goal-weight values. **(A)** Simulation with *δ* = 1. **(B)** Simulation with *δ* = 2. **(C)** Simulation with *δ* = 3. **(D)** Simulation with *δ* = 4. Each parameter setting was simulated using 32 agents. The *habit weight* P was fixed at 1, and the *context stability bias φ* was fixed at 0.5. The layout follows that of Figure 3.

**Figure 13:**
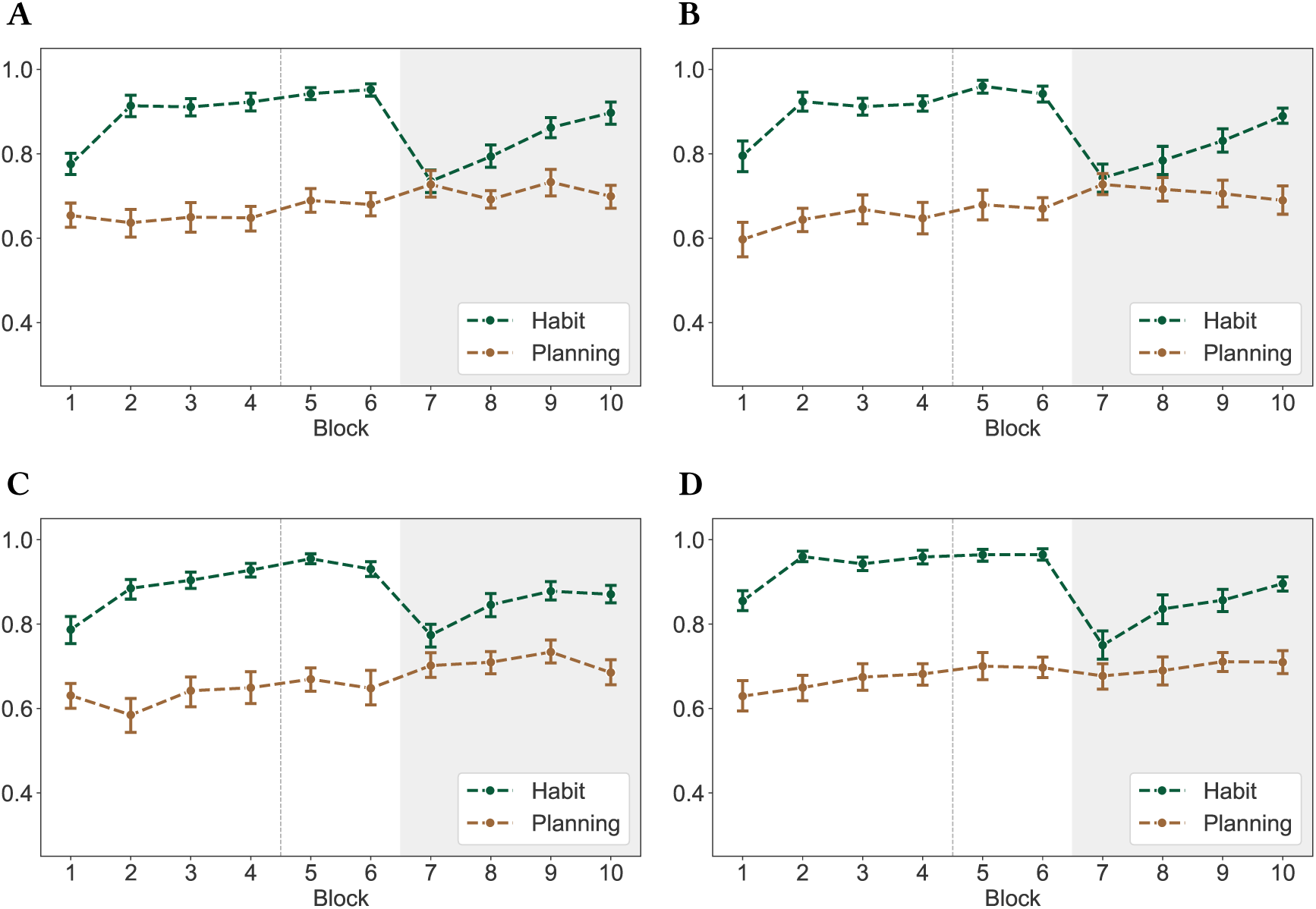
Optimal-choice rates across context-stability-bias values. **(A)** Simulation with *φ* = 0.25. **(B)** Simulation with *φ* = 0.50. **(C)** Simulation with *φ* = 0.75. **(D)** Simulation with *φ* = 1.00. Each parameter setting was simulated using 32 agents. The *goal weight δ* was fixed at 2, and the *habit weight* P was fixed at 1. The aggregated optimal-choice rates varied little across values of the context stability bias, likely because context inference was driven primarily by the context cue (background color) presented on each trial. The layout follows that of Figure 3.

**Figure 14:**
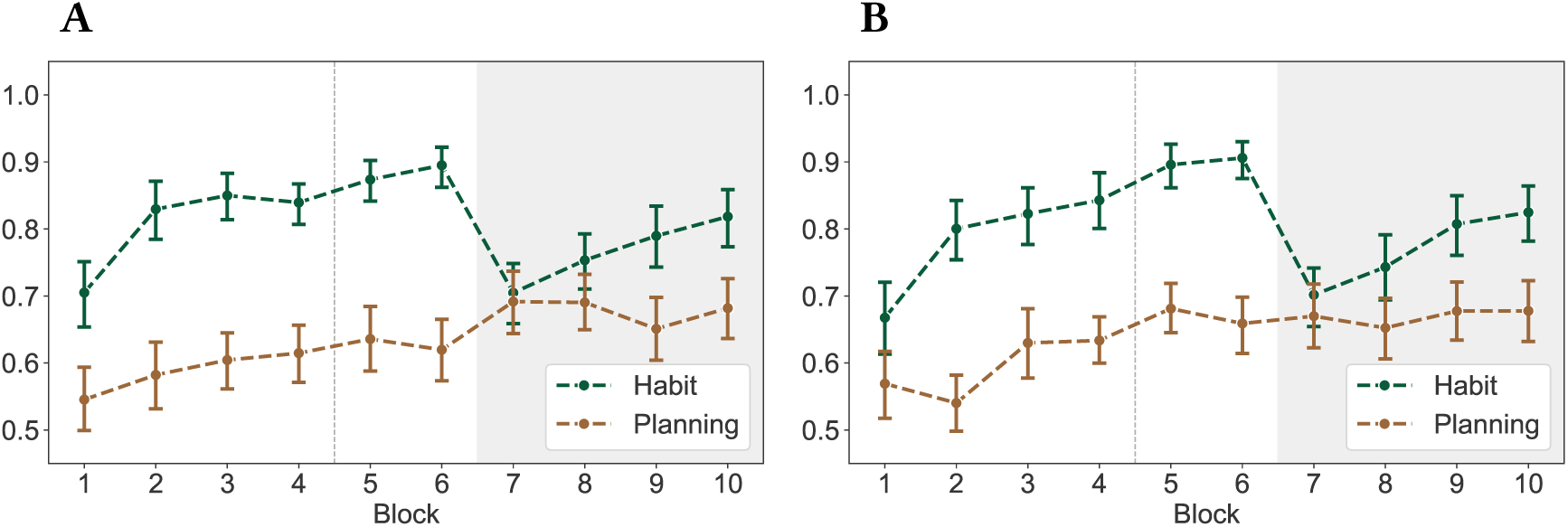
Posterior predictive checks of optimal-choice rates for the BCC-2P and BCC-3P models. **(A)** Posterior predictions for the BCC-2P model. **(B)** Posterior predictions for the BCC-3P model. Simulations were generated using the participantlevel posterior means of the inferred parameters. For the BCC-2P model, the context stability bias parameter *φ* was fixed at 0.46. The layout follows that of Figure 3.

**Figure 15:**
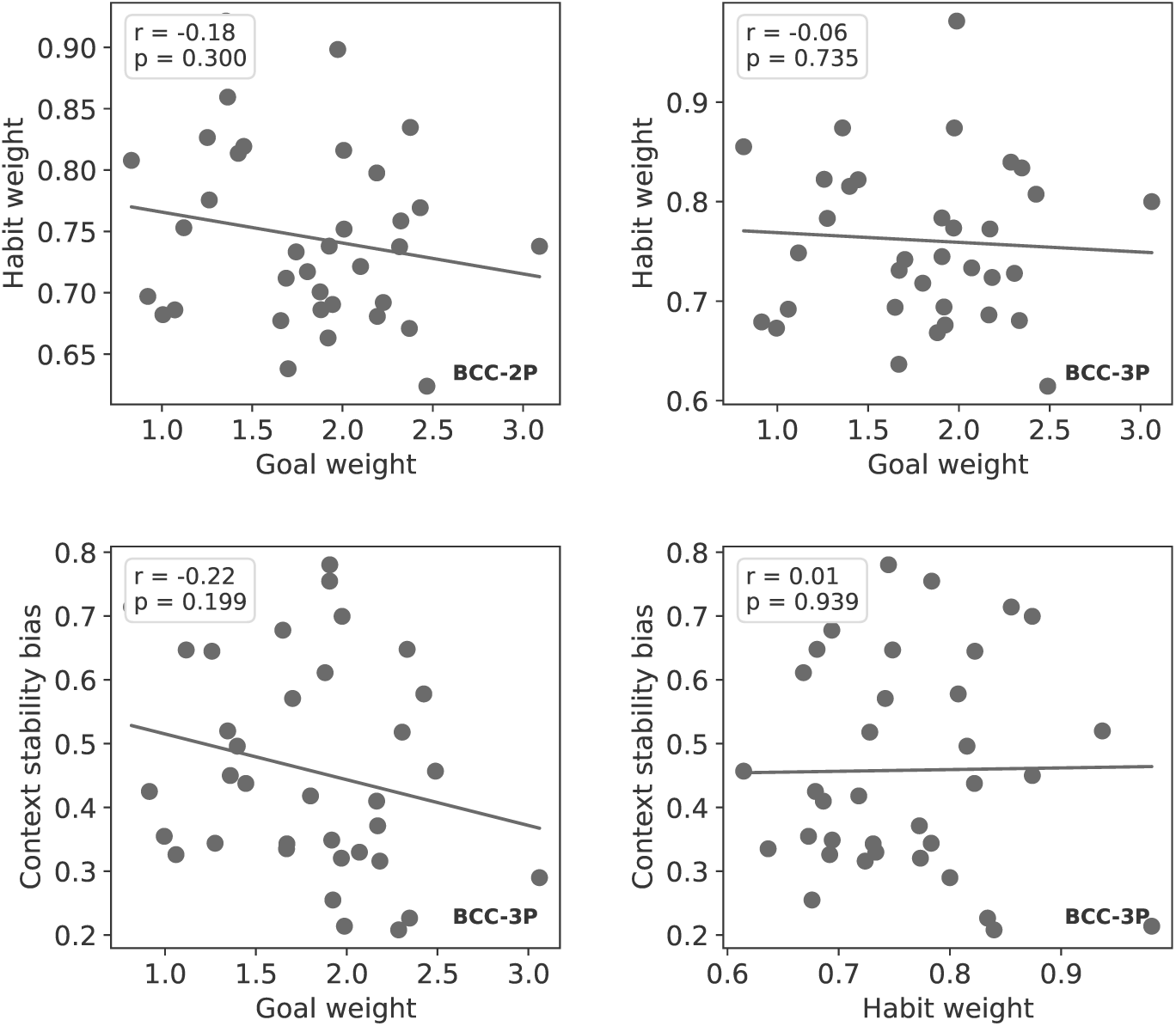
Correlations for inferred parameter values for the BCC-2P and BCC-3P models. Pearson correlation coefficients *r* and significance values *p* were calculated using the the participant-level posterior means of each parameter.

#### A.5 Miscellaneous

**Interpretation of Mixed-Effects Model Notation** Mixed-effects models are reported using notation based on the formula syntax of the lme4 package in R. For example, the model predicting optimal choices during acquisition was specified as

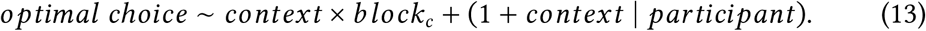

In the fixed-effects part of the model, the expression *context* × *block_c_* includes the two main effects and their interaction:

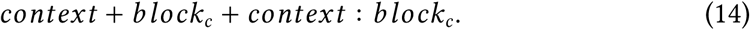

The term (1 + *context* ∣ *participant*) specifies a by-participant random intercept and a by-participant random slope for context. The vertical bar identifies participant as the grouping factor. The random intercept and random context slope are allowed to be correlated. For a formal specification, let *Y_ij_* denote the binary accuracy outcome on trial *j* for participant *i*, with *Y_ij_* = 1 indicating an optimal choice and *Y_ij_* = 0 indicating a non-optimal choice. The outcome is assumed to follow a Bernoulli distribution:

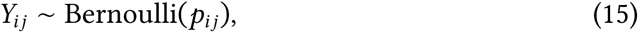

where *p_ij_* = *P*(*Y_ij_* = 1) is the probability of an optimal choice. The linear predictor is related to this probability through the logit link function:

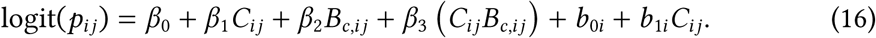

Here, *C_ij_* denotes the context predictor and *B_c_*_,*ij*_ denotes centered block. With planning as the reference category, *C_ij_* = 0 for planning trials and *C_ij_* = 1 for habit trials. The model parameters have the following interpretations:

- *β*_0_ is the population-level intercept. It represents the expected log-odds of an optimal choice in the planning context when centered block equals zero.
- *β*_1_ is the population-level context effect. It represents the habit-versus-planning difference when centered block equals zero.
- *β*_2_ is the population-level effect of block in the planning context.
- *β*_3_ is the context-by-block interaction. It represents the difference between the block effect in the habit context and the block effect in the planning context.
- *b*_0*i*_ is participant *i*’s deviation from the population-level intercept.
- *b*_1*i*_ is participant *i*’s deviation from the population-level context effect.

All mixed-effects models were fitted using lme4. Random-effects structures were specified using single-bar notation, allowing correlations between random intercepts and slopes when both were included.

